# Chromatin remodeler *Arid1a* regulates subplate neuron identity and wiring of cortical connectivity

**DOI:** 10.1101/2020.12.14.422645

**Authors:** Daniel Z. Doyle, Mandy M. Lam, Adel Qalieh, Yaman Qalieh, Alice Sorel, Owen H. Funk, Kenneth Y. Kwan

**Author notes:** Correspondence should be addressed to K.Y.K.

## Abstract

Loss-of-function mutations in chromatin remodeler gene *ARID1A* are a cause of Coffin-Siris syndrome, a developmental disorder characterized by dysgenesis of corpus callosum. Here, we characterize *Arid1a* function during cortical development and find unexpectedly selective roles for *Arid1a* in subplate neurons. Subplate neurons (SPNs), strategically positioned at the interface of cortical grey and white matter, orchestrate multiple developmental processes indispensable for neural circuit wiring. We find that pan-cortical deletion of *Arid1a* leads to extensive mistargeting of intracortical axons and agenesis of corpus callosum. Sparse *Arid1a* deletion, however, does not autonomously misroute callosal axons, implicating non-cell autonomous *Arid1a* functions in axon guidance. Supporting this possibility, the ascending axons of thalamocortical neurons, which are not autonomously affected by cortical *Arid1a* deletion, are also disrupted in their pathfinding into cortex and innervation of whisker barrels. Coincident with these miswiring phenotypes, which are reminiscent of subplate ablation, we unbiasedly find a selective loss of SPN gene expression following *Arid1a* deletion. In addition, multiple characteristics of SPNs crucial to their wiring functions, including subplate organization, subplate-thalamocortical axon co-fasciculation (“handshake”), and extracellular matrix, are severely disrupted. To empirically test *Arid1a* sufficiency in subplate, we generate a cortical plate deletion of *Arid1a* that spares SPNs. In this model, subplate *Arid1a* expression is sufficient for subplate-thalamocortical axon co-fasciculation and extracellular matrix assembly. Consistent with these wiring functions, subplate *Arid1a* sufficiently enables normal callosum formation, thalamocortical axon targeting, and whisker barrel development. Thus, *Arid1a* is a multifunctional regulator of subplate-dependent guidance mechanisms essential to cortical circuit wiring.

**Significance:** The cognitive, perceptive, and motor capabilities of the mammalian cerebral cortex depend on assembly of circuit connectivity during development. Subplate neurons, strategically located at the junction of grey and white matter, orchestrate the wiring of cortical circuits. Using a new approach to study gene necessity and sufficiency in subplate neurons, we uncover an essential role for chromatin remodeler *Arid1a* in subplate neuron gene expression and axon guidance functions. Cortical deletion of *Arid1a* disrupts subplate-dependent formation of corpus callosum, targeting of thalamocortical axons, and development of sensory maps. Together, our study identifies *Arid1a* as a central regulator of subplate-dependent axon pathfinding, establishes subplate function as essential to callosum development, and highlights non-cell autonomous mechanisms in neural circuit formation and disorders thereof.

## Introduction

The subplate is a transient layer of the fetal cerebral cortex essential to the developmental wiring of cortical circuits (1-11). During neurogenesis, cortical neural progenitor cells (NPCs) generate excitatory neurons following an orderly temporal progression, successively giving rise to subplate neurons (SPNs), then deep layer neurons, then upper layer neurons (12, 13). As the first neurons generated from embryonic cortex, SPNs establish emerging axon tracts and form the earliest synapses (1-4, 14-17). Importantly, SPNs, which are strategically positioned at the interface between post-migratory neurons and the developing white matter, serve non-cell autonomous wiring functions in the formation of cortical circuits. Experimental subplate ablation during fetal development leads to misrouting of thalamocortical axons (18-20) and disrupts formation of sensory maps (21, 22), and perturbed subplate function has been hypothesized to contribute to circuitry defects in disorders of brain development (23-25). Mechanistically, SPNs non-cell autonomously mediate circuit wiring at least in part by extending the earliest cortical descending axons, which interact with ascending thalamocortical axons during pathfinding (as posited by the “handshake hypothesis”) (26, 27) and contribute to their crossing of the pallial-subpallial boundary (PSB) (28, 29). SPNs also secrete extracellular matrix components that support axon guidance (4, 30). In addition, SPNs are required for early oscillatory activity (8, 14). In postnatal ages, some SPNs undergo programmed cell death (31), thereby serving a transient role in cortical circuit development.

Despite the central position of SPNs in orchestrating cortical connectivities, the molecular determinants of subplate wiring functions have remained largely elusive. Previous studies have focused on genes selectively expressed in SPNs (24, 32). These important studies illuminated the genetic bases of SPN specification, migration, and axon development (33-39). The severe axon misrouting phenotypes of subplate ablation (18-20), however, are not broadly recapitulated in these genetic mutants. And the mechanisms underpinning the axon guidance functions of subplate have remained largely mysterious. Here, by cell type-specific dissection of gene function, we identify *Arid1a* as a key regulator of multiple subplate-dependent axon guidance mechanisms indispensable for cortical circuit wiring.

*Arid1a* (*Baf250a*) encodes a subunit of the Brg/Brahma-associated factors (BAF, or mammalian SWI/SNF) ATP-dependent chromatin remodeling complex that mobilizes nucleosomes along DNA, thereby mediating processes such as transcriptional regulation and DNA repair. Human genetic studies have identified loss-of-function mutations in *ARID1A* in intellectual disability, autism spectrum disorder, and Coffin-Siris syndrome, a developmental disorder characterized by callosal dysgenesis (40). Similar to some other chromatin remodeling complexes, the BAF complex is broadly present in cells and organs (41). Widely-expressed chromatin remodelers, however, can function cell type-dependently (42). For BAF, cell type-dependent subunit compositions of the complex have been shown to play diverse roles in embryonic stem cells, NPCs, and neurons (43). *Arid1a*, however, has not been found to be incorporated into BAF complex in a cell type-specific way and its potential context-dependent roles remain to be fully explored.

Here, we leverage a conditional allele to cell type-specifically manipulate *Arid1a* function. We find that pan-cortical deletion of *Arid1a* leads to extensive mistargeting of intracortical axons and agenesis of corpus callosum. Surprisingly, unlike pan-cortical deletion, sparse *Arid1a* deletion does not cell autonomously misroute callosal axons, suggesting that axon mistargeting is a non-cell autonomous consequence of pan-cortical *Arid1a* deletion. Supporting this possibility, the axons of thalamocortical neurons, which are not autonomously affected by cortical *Arid1a* deletion, are also strikingly disrupted in their pathfinding into cortex and in their innervation of whisker barrels. *Arid1a* thus plays essential, non-cell autonomous roles in the development of multiple cortical axon tracts. At the transcriptomic level, we unbiasedly find a selective loss of SPN gene expression following *Arid1a* deletion, thus identifying subplate as a potential substrate of *Arid1a* phenotypes. Consistent with this, *Arid1a* axon misrouting defects are highly reminiscent of subplate ablation (18-20). Furthermore, multiple characteristics of SPNs crucial to their circuit wiring functions, including subplate organization and extracellular matrix, are disrupted following *Arid1a* deletion. Importantly, descending subplate axons are severely attenuated, abrogating their co-fasciculation with ascending thalamocortical axons. This “handshake” interaction with subplate axons is essential to thalamocortical axon pathfinding and whisker barrel formation (26, 27), both of which are disrupted by *Arid1a* deletion. Thus, we find a necessity for *Arid1a* in orchestrating distinct aspects of subplate circuit wiring functions. To empirically test *Arid1a* sufficiency in SPNs, we use a genetic approach to generate a cortical plate deletion of *Arid1a* that spares SPNs. In this model, we find that *Arid1a* expression in SPNs is sufficient to support subplate organization, “handshake” with thalamocortical axons, and extracellular matrix. Consistent with these wiring functions, subplate *Arid1a* expression sufficiently enables normal thalamocortical axon targeting, whisker barrel development, and callosum formation. Together, our study identifies *Arid1a* as a central regulator of subplate-dependent axon pathfinding, unequivocally establishes subplate function as essential to callosal development, and highlights non-cell autonomous mechanisms in circuit development and disorders thereof.

## Results

### Mistargeting of intracortical, but not corticofugal, axons following *Arid1a* deletion from NPCs

We analyzed ARID1A protein in the developing mouse cortex by immunostaining and found broad ARID1A expression in ventricular zone (VZ) and subventricular zone (SVZ) NPCs and cortical plate (CP) neurons from embryonic day (E)11.5 to postnatal day (P)0 (**Fig. 1*A***). We found complete colocalization of ARID1A with DAPI DNA staining at E14.5 (***SI Appendix*, Fig. S1*A***), which is consistent with ubiquitous ARID1A expression during cortical development. Constitutive deletion of *Arid1a* leads to lethality on E6.5 in mice (44). We therefore leveraged a conditional *Arid1a* allele (44) to investigate *Arid1a* function during brain development.

**Fig. 1.**
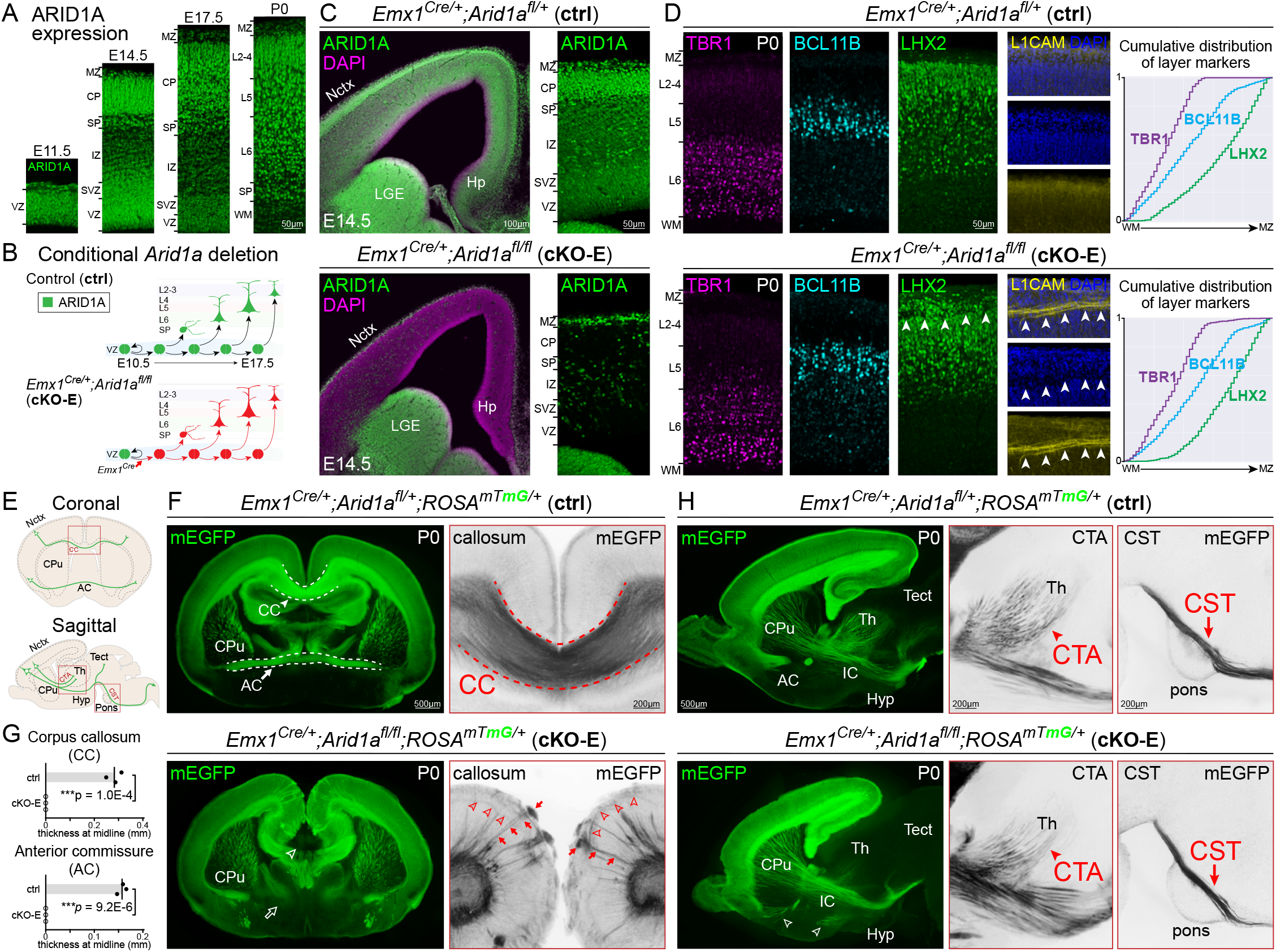
Tract-dependent misrouting of cortical axons following conditional *Arid1a* deletion. (*A*) ARID1A (green) immunostaining on coronal embryonic day (E)11.5, E14.5, E17.5, and postnatal day (P)0 brain sections revealed widespread ARID1A expression during cortical development. (*B*) Schematic illustration of conditional *Arid1a* deletion using *Emx1*^*Cre*^, which mediates recombination in cortical NPCs at E10.5, near the onset of neurogenesis. (*C*) ARID1A (green) and DAPI (magenta) staining of coronal E14.5 *Emx1*^*Cre/+*^*;Arid1a*^*fl/fl*^ (cKO-E) brain sections revealed loss of ARID1A from VZ and SVZ NPCs and CP neurons derived from *Emx1* lineage. ARID1A expression was unaffected in ventral forebrain. (*D*) Layer marker immunostaining on coronal P0 brain sections. TBR1+ (L6, magenta), BCL11B+ (L5, cyan), and LHX2+ (L2-5, green) neurons were correctly ordered in cKO-E. Analysis of cumulative distribution of layer marker-expressing neurons through thickness of cortex from white matter (WM) to marginal zone (MZ) revealed no disruption in cortical lamination in cKO-E (n=3 animals). In all analyzed cKO-E brains (3/3 animals), but none of the littermate control (ctrl) brains (0/3 animals), a stereotyped gap (arrowheads) in the upper cortical layers was observed in LHX2 and DAPI (blue) staining. This gap contained misrouted L1CAM+ (yellow) axons. (*E*) Schematic illustration of cortical axon tracts on coronal and sagittal brain sections. (*F*) Coronal sections of P0 ctrl and cKO-E brains. Membrane EGFP (mEGFP, green) was expressed Cre-dependently from *ROSA*^*mTmG*^, enabling visualization of cortical axons. Agenesis of corpus callosum (open arrowhead) was observed in cKO-E (n=3/3 animals). The anterior commissure also failed to form (open arrow). The cKO-E cortex was characterized by widespread axon misrouting (n=3/3 animals), including radially-directed axons extending to the pia (red arrows) and tangentially-directed axons travelling across the upper layers (red arrowheads). (*G*) Quantification of corpus callosum and anterior commissure thickness at midline (data are mean, two-tailed unpaired *t* test, n=3 animals). (*H*) Sagittal sections of P0 ctrl and cKO-E brains. In cKO-E, corticofugal axons innervated internal capsule (IC) without defect. Corticothalamic axons (CTA, red arrowheads), corticotectal axons, and corticospinal tract axons (CST, red arrows) were qualitatively reduced, but followed the normal trajectories without misrouting defects in cKO-E. Axons from anterior commissure (AC) were misrouted to hypothalamus (Hyp, open arrowheads). VZ, ventricular zone; SVZ, subventricular zone; PP, preplate; IZ, intermediate zone; SP, subplate; CP, cortical plate; MZ, marginal zone; Ln, layer n; WM, white matter; Nctx, neocortex; Hp, hippocampus; LGE, lateral ganglionic eminence; CC, corpus callosum; AC, anterior commissure; CPu, caudate putamen; CTA, corticothalamic axons; CST, corticospinal tract; Th, thalamus; Tect, tectum; Hyp, hypothalamus

To delete *Arid1a* from developing cortex, we used *Emx1*^*Cre*^, which expresses Cre recombinase in cortical NPCs starting at ∼E10.5, near the onset of cortical neurogenesis (**Fig. 1*B***) (45). Deletion of *Arid1a* was validated in *Emx1*^*Cre/+*^;*Arid1a*^*fl/fl*^ conditional mutants (**cKO-E**) by ARID1A immunostaining, which revealed loss of ARID1A from cortical NPCs and neurons of the *Emx1* lineage (**Fig. 1*C***). Mutant cKO-E mice were born at Mendelian ratio, had typical lifespan, and were fertile as adults. At P0, the size of cKO-E cortex was not significantly different from control (ctrl) littermates (***SI Appendix*, Fig. S1 *B* and *C***).

To assess neocortical layers, we analyzed laminar markers at P0 (**Fig. 1*D***). We found largely normal cortical lamination in cKO-E; TBR1+ layer (L)6, BCL11B+ (CTIP2+) L5, and LHX2+ L2-5 neurons were properly ordered and not robustly different from control based on marker quantification (***SI Appendix*, Fig. S1*D***) and analysis of cumulative distribution of layer markers (**Fig. 1*D***). However, a stereotyped gap in marker and DAPI staining was present in upper cortical layers (L2-4) of all analyzed cKO-E brains (arrowheads, **Fig. 1*D***, 3/3 animals) but absent from controls (0/3 animals). This gap was characterized by aberrant L1CAM-immunostained axons (arrowheads, **Fig. 1*D***), suggesting the presence of misrouted axons in cKO-E cortex.

We next examined potential changes in axonal projections using the Cre-dependent fluorescent reporter allele *ROSA*^*mTmG*^ (46). Following *Emx1*^*Cre*^ recombination, membrane EGFP (mEGFP) was expressed from *ROSA*^*mTmG*^ in all cortical excitatory neurons, enabling visualization of their axon projections (schematized in **Fig. 1*E***). In P0 control, mEGFP expression revealed corpus callosum (CC), which connected the cortical hemispheres, and anterior commissure (AC), which connected lateral regions (**Fig. 1*F***). In cKO-E, we found corpus callosum agenesis and widespread misrouting of cortical axons, including radially-directed axons towards the pia (arrows) and tangentially-directed axons through the upper layers (arrowheads, **Fig. 1*F***). These misrouting phenotypes were confirmed by L1CAM immunostaining (***SI Appendix*, Fig. S1*E***). Anterior commissure axons were also mistargeted and unable to cross the midline (**Fig. 1*F*** and ***SI Appendix*, Fig. S1*F***). Analysis of callosal and commissural thickness revealed complete loss of these tracts at the midline in cKO-E (**Fig. 1*G***). Axon misrouting was also present in hippocampus, and accompanied hippocampal hypoplasia and disorganization (***SI Appendix*, Fig. S1*G***).

In contrast to intracortical axons, corticofugal tracts (i.e. corticothalamic, corticotectal, and corticospinal) were not characterized by gross misrouting deficits in cKO-E (**Fig. 1*H***). These tracts were qualitatively reduced in strength but followed normal trajectories out of the cortex, through internal capsule (IC), and innervated their respective targets in dorsal thalamus, tectum, and medulla. Together, our axonal analyses revealed widespread mistargeting of intracortical, but not corticofugal axon tracts following *Arid1a* deletion, implicating a tract-dependent role for *Arid1a* in cortical circuits.

### Non-cell autonomous misrouting of thalamocortical axons following *Arid1a* deletion

We next analyzed thalamocortical axons (schematized in **Fig. 2*A***). In cKO-E, thalamocortical neurons were not autonomously affected by cortical *Arid1a* deletion; however, the target of these axons, the cortex, was broadly affected by *Emx1*^*Cre*^. We used cytochrome oxidase (CO) histochemistry on flattened P7 cortex to visualize whisker barrels, a major target of thalamocortical axons (TCAs). In P7 control, CO staining revealed discrete, stereotyped, and organized whisker barrels in primary somatosensory cortex (**Fig. 2*B*** and ***SI Appendix*, S2*A***). In cKO-E, barrel formation was severely disrupted; many barrels were missing and the remaining barrels were distorted and disorganized, suggesting a deficit in thalamocortical axon targeting. These defects were confirmed by immunostaining for NTNG1 (Netrin G1), a marker of thalamocortical axons (**Fig. 2*B***). In addition, analysis of P0 cKO-E revealed that NTNG1+ axons deviated from their normal trajectory in white matter (WM) (arrowheads, **Fig. 2*C***) and became markedly misrouted, including into tangential bundles through upper cortical layers and toward the midline. Interestingly, these thalamocortical axons largely did not contribute to the radially-directed aberrant axons in cKO-E, which were revealed by pan-axonal marker L1CAM (arrows, **Fig. 2*C***) and *ROSA*^*mTmG*^ expression (arrows, ***SI Appendix*, Fig. S2*B***).

**Fig. 2.**
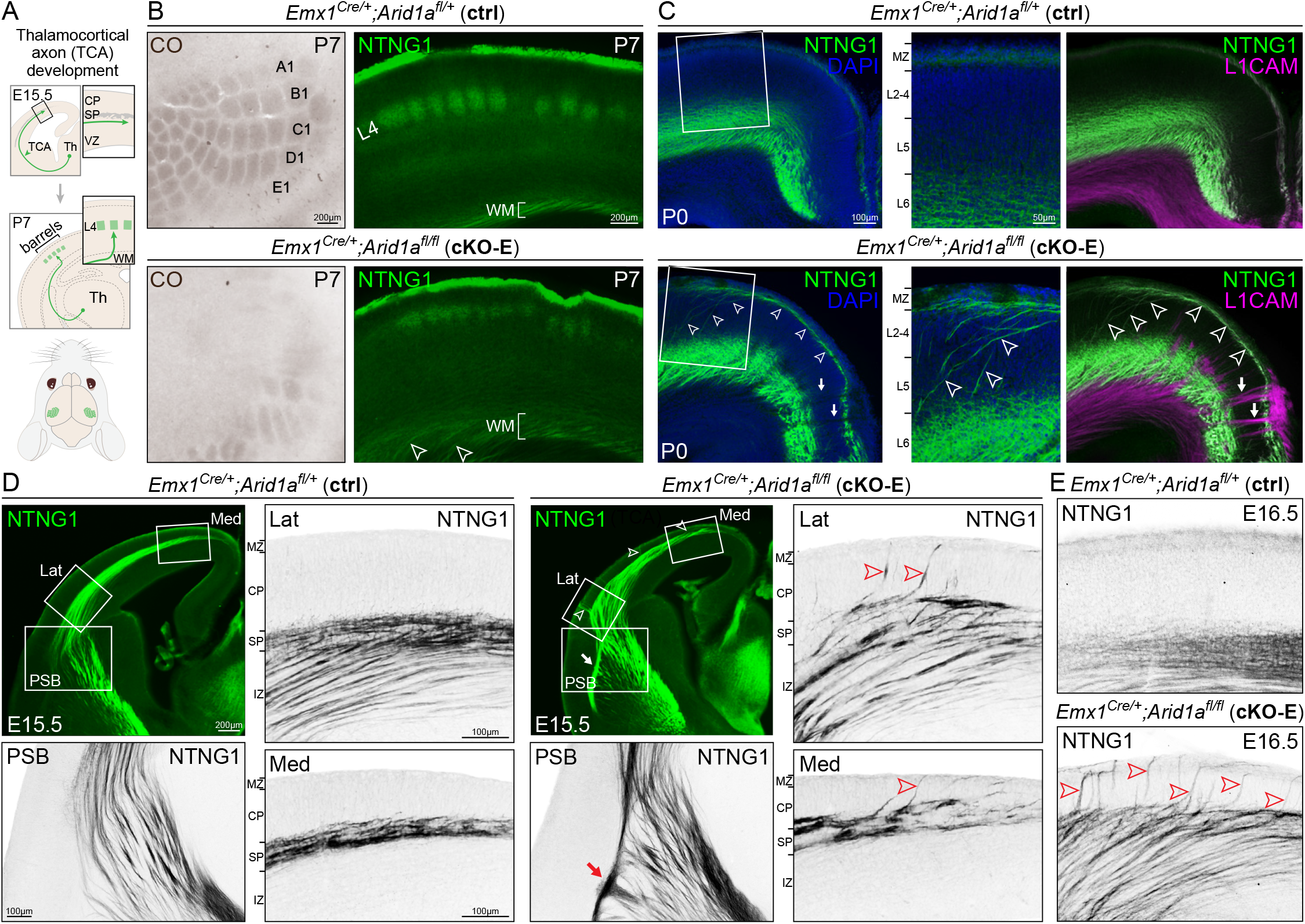
Non-cell autonomous disruption of thalamocortical axon pathfinding following *Arid1a* deletion. (*A*) Schematic illustration of thalamocortical axon (TCA) development. (*B*) Whisker barrels in P7 ctrl and cKO-E primary somatosensory cortex were visualized by cytochrome oxidase staining (CO, brown) on flattened cortices and NTNG1 immunostaining (green) on coronal sections. In cKO-E, barrel formation was severely disrupted (n=4/4 animals). Many barrels were missing and the remaining barrels were distorted or disorganized. NTNG1+ thalamocortical axons were defasciculated in cortical white matter (open arrowheads) in cKO-E. (*C*) NTNG1 immunostaining (green) on coronal sections of P0 ctrl and cKO-E brains. In ctrl, NTNG1+ thalamocortical axons were present in white matter and L6, as well as marginal zone (MZ). In cKO-E, NTNG1+ thalamocortical axons were markedly misrouted, extending dorsally from white matter through the cortical layers (arrowheads). These aberrant axons then travelled tangentially across the upper layers and toward the midline. NTNG1+ thalamocortical axons did not contribute to the abnormal radially-directed axon bundles labeled by L1CAM (magenta, arrows) in cKO-E. (*D*) Analysis of thalamocortical axon development in E15.5 ctrl and cKO-E cortex. In ctrl, NTNG1+ thalamocortical axons extended across the pallial-subpallial boundary (PSB) and were paused within the subplate (SP) during the embryonic “waiting” period. In cKO-E, NTNG1+ thalamocortical axons did not cross the PSB along the normal trajectory (n=3/3 animals). They formed an aberrant bundle of axons parallel to the PSB (red arrow) and entered the cortex via an abnormal medial path. Notably, NTNG1+ thalamocortical axons prematurely invaded the cortical plate (CP) in both lateral (Lat) and medial (Med) cortex (red arrowheads) (n=3/3 animals). The trajectory of these axons was similar to that of the misrouted thalamocortical axons in the P0 cKO-E cortex (arrowheads in *C*). (*E*) Analysis of E16.5 ctrl and cKO-E cortex revealed an abundance of NTNG1+ thalamocortical axons prematurely invading the cortical plate (red arrowheads).

Thalamocortical axons normally follow a precise developmental path. In control E15.5 cortex, NTNG1+ axons had crossed the pallial-subpallial border (PSB) and reached the subplate (SP). Consistent with the “waiting period” (4, 47, 48), thalamocortical axons at this age paused their ingrowth within the subplate and had not entered the cortical plate (CP, **Fig. 2*D***). Remarkably, in cKO-E, NTNG1+ axons prematurely invaded the cortical plate as aberrant bundles of axons (arrowheads, **Fig. 2*D***). By E16.5, we found an abundance of thalamocortical axons prematurely invading cortical plate (**Fig. 2*E***). This bypassing of the waiting period by thalamocortical axons is reminiscent of subplate ablation (18, 19). In addition, cKO-E NTNG1+ axons did not cross the PSB along the normal path. Instead, they formed an aberrant bundle running parallel to the PSB (arrow, **Fig. 2*D*** and ***SI Appendix*, Fig. S2*C***) and ultimately entered the pallium via a narrow medial trajectory. Similar to bypassing the waiting period, disrupted PSB crossing of thalamocortical axons is also a known consequence of disrupted subplate function (28, 29). Together, misrouting of thalamocortical axons following cortex-specific *Arid1a* deletion in cKO-E revealed non-cell autonomous *Arid1a* functions in axon guidance.

### Correct callosal axon targeting following sparse deletion of *Arid1a*

To determine whether the intracortical axon misrouting in cKO-E was also a result of disrupted non-cell autonomous *Arid1a* function, we sought to sparsely delete *Arid1a* from cortical neurons by *in utero* electroporation (IUE). We generated a self-excising, self-reporting *Cre*-recombinase construct (CAG-sxiCre-EGFP) that would express Cre, and upon Cre recombination, simultaneously excise *Cre* and turn on EGFP expression (***SI Appendix*, Fig. S3*A***). The sxiCre construct was transfected into embryonic cortex by IUE (38) at E14.5 to target NPCs during genesis of upper-layer neurons, which form the majority of callosal axons. Transfected brains were analyzed at P0. To directly compare the effects of sparse versus widespread loss of *Arid1a*, we used control, cKO-E (pan-cortical), and *Arid1a*^*fl/fl*^ (sparse by Cre transfection) littermates for IUE. The presence of ARID1A in EGFP+ electroporated cells was determined by immunostaining; sxiCre had a 97.67% deletion efficacy in *Arid1a*^*fl/fl*^ (***SI Appendix*, Fig. S3*B***). In P0 control, electroporated neurons migrated to the upper cortical layers and projected EGFP+ axons across the corpus callosum (**Fig. 3 *A-C***). Following broad, genetic deletion of *Arid1a* in cKO-E, electroporated neurons migrated to the upper layers, although their positioning was qualitatively less organized and potentially affected by misrouted axons (**Fig. 1*D***). In cKO-E, axons originating from transfected neurons failed to project into corpus callosum (**Fig. 3 *D-F***) and contributed to aberrant radially-directed bundles (***SI Appendix*, Fig. S3*C***). Following sparse *Arid1a* deletion by sxiCre IUE in *Arid1a*^*fl/fl*^ mice, electorporated neurons migrated to upper layers without deficit (**Fig. 3 *G*** and ***H***). Remarkably, despite absence of ARID1A, these neurons abundantly extended EGFP+ axons into the corpus callosum and correctly targeted contralateral cortex in a manner indistinguishable from control (**Fig. 3*I***). Together, our results provided strong support that the callosal and thalamocortical axon pathfinding defects in cKO-E were non-cell autonomous consequences of broad cortical *Arid1a* deletion.

**Fig. 3.**
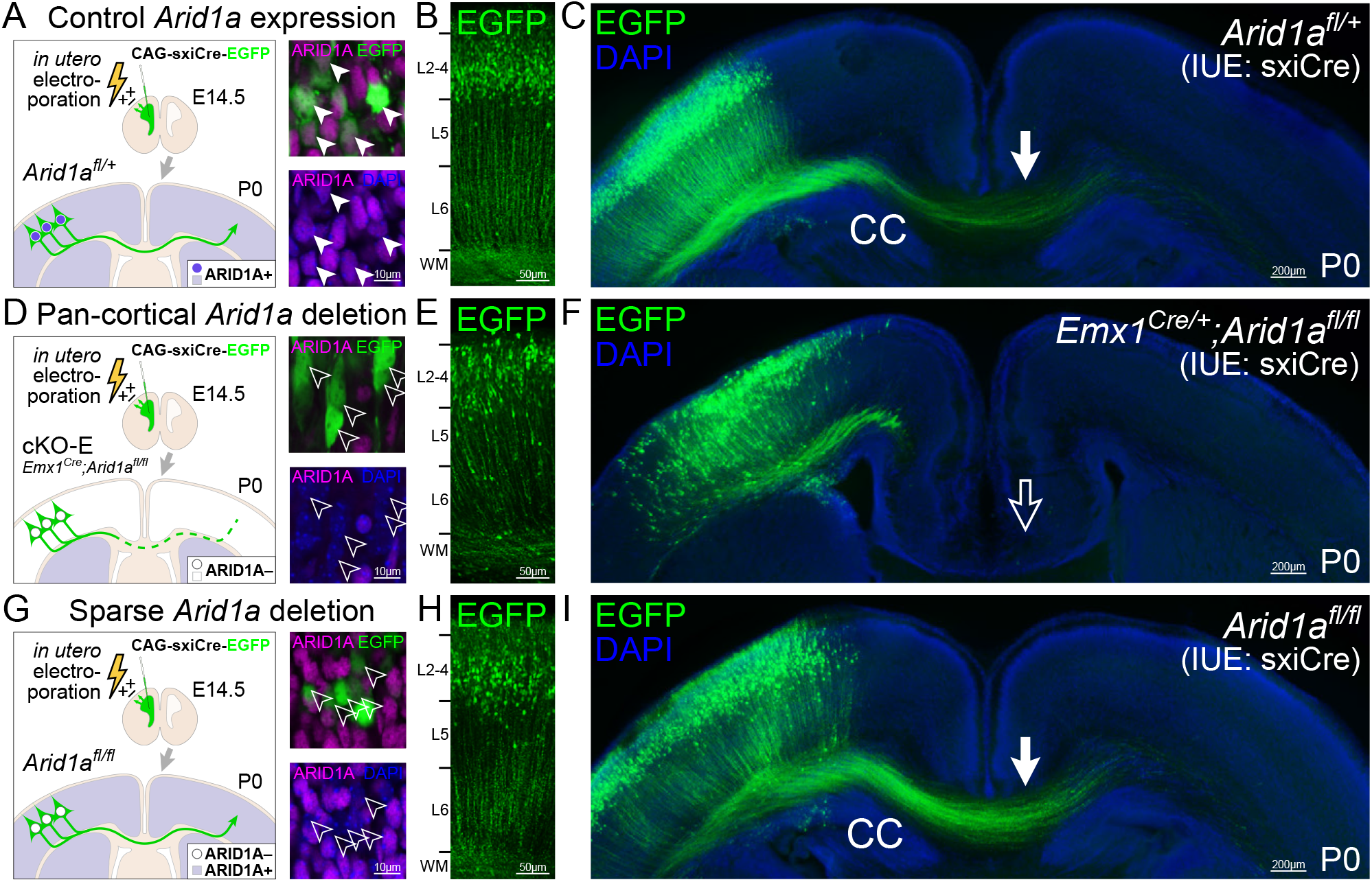
Correct callosal axon targeting following sparse deletion of *Arid1a*. A self-excising Cre expression EGFP reporter construct (CAG-sxiCre-EGFP or sxiCre) was transfected into dorsal cortical NPCs of *Arid1a*^*fl/+*^ (control, *A*), *Emx1*^*Cre/+*^*;Arid1a*^*fl/fl*^ (cKO-E, *D*), and *Arid1a*^*fl/fl*^ (without genetic Cre, *G*) using *in utero* electroporation at E14.5. At P0, ARID1A expression (magenta) was analyzed in EGFP+ transfected cells by immunostaining. ARID1A was present in transfected control EGFP+ cells (solid arrowheads, a), but was lost following pan-cortical genetic *Arid1a* deletion (cKO-E, open arrowheads, *D*) or sparse *Arid1a* deletion (*Arid1a*^*fl/fl*^, open arrowheads, *G*). EGFP+ cells migrated to the upper cortical layers in each condition (*B, E, H*). EGFP+ axons innervated the corpus callosum (CC) in control (solid arrow, *C*), but failed to do so following broad *Arid1a* deletion in cKO-E (open arrow, *F*). Remarkably, sparse deletion of *Arid1a* from *Arid1a*^*fl/fl*^ EGFP+ cells did not disrupt their innervation of the corpus callosum (solid arrow, *I*). Loss of ARID1A expression from these cells (open arrowheads, *G*) following sparse *Arid1a* deletion did not cell autonomously cause a callosal axon misrouting defect.

### Selective disruption of subplate neuron gene expression following *Arid1a* deletion

To gain mechanistic insights into the axon guidance roles of *Arid1a*, we explored its molecular functions. As a chromatin remodeler, ARID1A mediates transcriptional regulation (49, 50). We thus analyzed the transcriptomes of cKO-E and littermate control cortices at E15.5, a developmental stage when cortical and thalamocortical axons undergo pathfinding (51). We generated libraries for unique molecular identifier (UMI) RNA-seq using Click-seq (52-54) from E15.5 cKO-E and littermate control cortices (ctrl: n=6, cKO-E: n=6 animals). The wide dynamic range of UMI RNA-seq was confirmed by spike-in ERCC standards (***SI Appendix*, Fig. S4*A***). Differential gene expression was analyzed using edgeR (55), which revealed significant differential expression of 103 annotated genes in cKO-E compared to control with a stringent false discovery rate (FDR) of <0.01 (**Fig. 4*A***). Gene expression changes were quantitatively validated by droplet digital (dd)RT-PCR for two downregulated genes (*Tle4, Zfpm2*) and two genes without significant changes (*Lhx2, Tbr1*), and by immunostaining for TLE4 (***SI Appendix*, Fig. S4*B***).

**Fig. 4.**
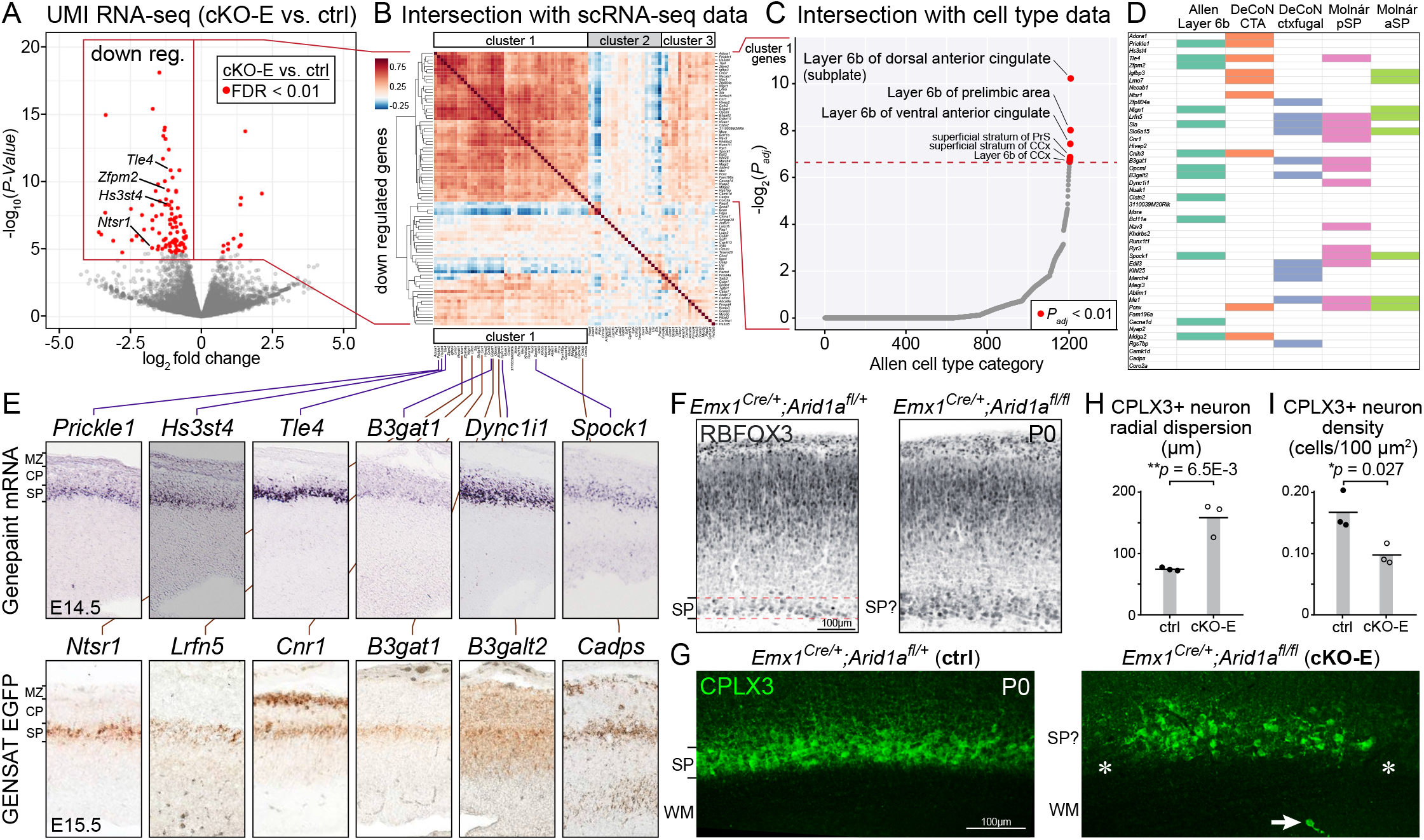
Selective disruption of subplate neuron gene expression following *Arid1a* deletion. (*A*) Volcano plot of unique molecular identifier (UMI) RNA-seq data comparing E15.5 cortex of cKO-E (n=6 animals) to ctrl littermates (n=6 animals). For each gene, *P*-value was calculated with likelihood ratio tests and false discovery rate (FDR) was calculated using the Benjamini-Hochberg procedure. Differentially expressed genes (FDR < 0.01) are indicated by red dots. Of the 103 differentially expressed genes, 91 were downregulated and 12 were upregulated in cKO-E. (*B*) Intersectional analysis of significantly downregulated genes with single cell (sc)RNA-seq data from wildtype embryonic forebrain (Yuzwa et al., 2017). Unsupervised hierarchical clustering revealed a cluster of 46 downregulated genes (cluster 1) that are highly co-expressed at the level of single cells, suggesting that they may be expressed from one cell type. (*C*) Intersectional analysis of the 46 genes in cluster 1 with a spatiotemporal gene expression dataset covering over 1200 brain subregions (Sunkin et al., 2013). Cluster 1 showed a significant overrepresentation of genes selectively expressed in cortical layer 6b. Layer 6b is alternative nomenclature for subplate. (*D*) Intersectional analyses with orthogonal datasets (Molyneaux et al., 2015; Oeschger et al. 2012, Sunkin et al., 2013) confirmed subplate expression of genes in cluster 1. (*E*) Subplate expression of cluster 1 genes was further confirmed by E14.5 *in situ* hybridization data from the Genepaint database and E15.5 EGFP transgene expression data from the GENSAT consortium (61, 62). (*F*) RBFOX3 immunostaining (black) on coronal sections of P0 ctrl revealed a distinct and organized subplate band positioned just beneath the cortical layers. In cKO-E, the subplate band was less clearly defined and not distinct from cortical layers. (*G*) CPLX3 immunostaining (green) on coronal sections of P0 ctrl and cKO-E brains. In ctrl, CPLX3+ subplate neurons (SPNs) were organized into a discrete, continuous band. In cKO-E, the band of CPLX3+ SPNs was more dispersed and characterized by gaps (asterisks). Some CPLX3+ cells were aberrantly positioned in white matter (WM, arrow). (*H, I*) Quantification of CPLX3+ SPN radial dispersion and density at P0 (data are mean, two-tailed unpaired *t* test, n=3 animals). pSP, posterior subplate; aSP, anterior subplate

*Arid1a* has been shown to increase or maintain transcriptional activity (50), which is consistent with our finding that a majority of differentially expressed genes was downregulated in cKO-E (91/103). To determine whether the axon defects in cKO-E may be associated with reduced expression of axon extension or axon guidance genes, we intersected the 91 downregulated genes with 253 axon guidance genes (GO:0007411) and 138 axon extension genes (GO:0048675). This revealed an overlap of one axon guidance gene (*Ablim1*) and one axon extension gene (*Myo5b*), which were validated by ddRT-PCR (***SI Appendix*, Fig. S4*C***). Neither group was significantly overrepresented (*P*_*hyper*_ = 0.34 and 0.36, respectively).

To determine whether gene expression from particular cortical cell types were preferentially affected in cKO-E, we performed intersectional analysis with single cell RNA-seq (scRNA-seq) data from wildtype embryonic cortex (56). Unsupervised hierarchical clustering of cKO-E downregulated genes based on scRNA-seq revealed a cluster with 46/91 genes (cluster 1) that are highly co-expressed at the level of single cells (**Fig. 4*B***), suggesting that these downregulated genes are normally expressed from one cell type. To validate this important finding, we performed intersectional analysis using an orthogonal dataset from wildtype E14.5 cortex (57), which revealed single-cell co-expression of 58/91 downregulated genes (cluster A, ***SI Appendix*, Fig. S4*D***). Remarkably, each of the 46 genes in cluster 1 was also represented in cluster A. This complete overlap provided high confidence that the downregulated genes in cKO-E reflected selective disruption of a single cell type.

To unbiasedly determine the identity of this cell type, we intersected the 46 cluster 1 genes with a rich spatiotemportal gene expression dataset covering over 1200 brain subregions (58) using Enrichr (59). This revealed a statistically significant enrichment of genes selectively expressed in cortical layer 6b, which is an alternative nomenclature for subplate (**Fig. 4*C***). We next intersected cluster 1 genes with additional datasets orthogonal to the discovery data. We found overrepresentation of cluster 1 genes in genes preferentially expressed in subplate based on microdissection (32) (**Fig. 4*D***). Intersection with a cell-type specific RNA-seq dataset (60) revealed overrepresentation of cluster 1 genes in “corticothalamic group 6” and “corticofugal group 11” (**Fig. 4*D*** and ***SI Appendix*, Fig. S4*E***). Although subplate neurons (SPNs) were not specifically annotated in this dataset, marker membership suggests that these two groups comprise SPNs. Notably, cluster 1 genes showed no significant overlap with other major cell types (e.g. callosal neurons, corticospinal neurons), suggesting selective dysregulation of SPN genes in cKO-E. Available E14.5 *in situ* hybridization data (Genepaint) (61) and E15.5 EGFP transgene expression data (GENSAT) (62) further supported that genes within cluster 1 are expressed in subplate (**Fig. 4*E***). Together, our analyses revealed selective disruption of SPN molecular identity following *Arid1a* deletion.

Consistent with disruption of SPN gene expression in cKO-E, anatomical development of SPNs was also altered. In P0 control, RBFOX3-labeled SPNs were organized in a distinct, tight band positioned beneath cortical plate (**Fig. 4*F***). In cKO-E, a discrete and organized subplate layer was absent; the subplate was intermingled with cortical plate. In control, SPN marker CPLX3 (63) specifically labeled neurons within a continuous subplate band (**Fig. 4*G***). In cKO-E, gaps were present in the subplate band (asterisks) and some CPLX3+ neurons were mispositioned in white matter. In addition, CPLX3+ SPNs were significantly more dispersed (**Fig. 4 *H*** and ***I***). Together, these data suggested that SPNs were present in cKO-E, but were anatomically disorganized and characterized by disrupted cell type-dependent gene expression.

### Altered organization and morphogenesis of embryonic subplate neurons following *Arid1a* deletion

The loss of SPN expression and non-cell autonomous axon phenotypes in cKO-E convergently suggested that *Arid1a* guidance functions may be centered on subplate. SPNs are indispensable for the wiring of cortical circuits. Subplate ablation misroutes cortical axons and causes thalamocortical axons to prematurely invade cortical plate in a manner reminiscent of cKO-E (**Fig. 2**) (18-20). Disrupted subplate function also leads to defects in thalamocortical axon crossing of the PSB and formation of cortical sensory maps (21, 22, 28, 29) similar to *Arid1a* deletion. We therefore characterized multiple SPN characteristics that contribute to their circuit wiring functions

Subplate mediates cortical axon pathfinding during embryonic ages. We thus assessed embryonic subplate organization by immunostaining of TUBB3 (TUJ1), a neuronal cytoskeleton marker that reveals the structure of embryonic cortical layers. In control E14.5 cortex, TUBB3+ processes were horizontally organized in intermediate zone (IZ). In cKO-E, however, TUBB3+ processes invaded cortical plate diagonally (**Fig. 5*A***). In control, analysis of MAP2, a somatodendritic marker, and NR4A2 (NURR1), a marker of SPNs and L6 neurons, revealed a horizontal organization of subplate as a distinct, continuous layer below cortical plate (**Fig. 5*B***). In cKO-E, SPNs showed abnormal clustering and cell-sparse gaps. Notably, MAP2-labeled dendrites aberrantly projected ventrally into intermediate zone (red arrowheads, **Fig. 5*B***). We further used the Tg(*Lpar1-EGFP*) transgenic allele, which expresses EGFP in a subset of SPNs (62, 64). In E16.5 and P0 cKO-E cortex, Lpar1-EGFP-labeled SPNs were characterized by cell-sparse gaps (asterisks) and some were aberrantly positioned in intermediate zone or white matter (arrows, **Fig. 5*C*** and ***SI Appendix*, Fig. S5*A***).

**Fig. 5.**
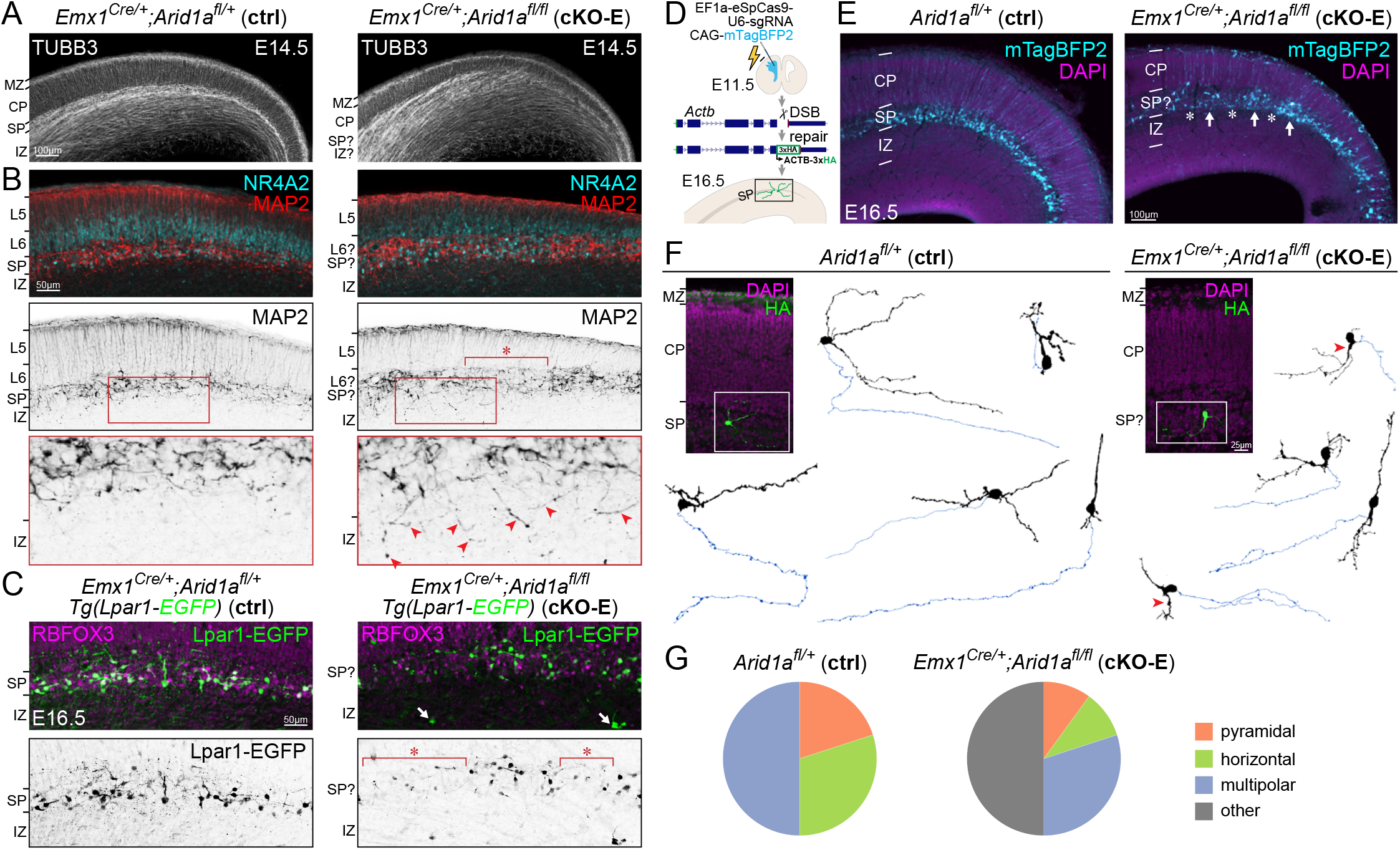
Disrupted subplate organization and SPN morphology following *Arid1a* deletion. (*A*) TUBB3 (TUJ1) immunostaining (white) on E14.5 ctrl and cKO-E sections. In ctrl, TUBB3+ processes were horizontally organized in intermediate zone (IZ) and subplate, and radially organized in cortical plate (CP). In cKO-E, TUBB3+ axons became defasciculated in intermediate zone, and invaded cortical plate diagonally. (*B*) MAP2 (red, black) and NR4A2 (cyan) immunostaining on E14.5 ctrl and cKO-E sections. In ctrl, MAP2+/NR4A2+ SPNs were organized within a clearly delineated layer below cortical plate. In cKO-E, SPNs were characterized by abnormal clustering and cell-sparse gaps (asterisk). In cKO-E, misoriented MAP2+ dendrites aberrantly projected ventrally into intermediate zone (red arrowheads, inset). (*C*) RBFOX3 (NEUN) immunostaining (magenta) on E16.5 ctrl and cKO-E brains carrying the *Lpar1-EGFP* transgene. In cKO-E, Lpar1-EGFP+ SPNs (green, black) were characterized by cell-sparse gaps (asterisks) and a few Lpar1-EGFP+ neurons were aberrantly positioned in the intermediate zone (arrows). (*D*) Schematic illustration of sparse SPN labeling by *in utero* genome editing. A DNA break was induced by CRISPR-Cas9 within the coding region of *Actb* near the C-terminus. A reporter repair template was designed such that correct DNA repair would lead to expression of ACTB-3xHA. CRISPR-Cas9, reporter repair, and mTagBFP2 expression constructs were co-transfected into cortical NPCs at E11.5 by *in utero* electroporation (IUE). Electroporated brains were analyzed at E16.5. (*E*) In electroporated E16.5 brains, mTagBFP2 (cyan) was successfully targeted to SPNs. In cKO-E, labeled SPNs showed disorganization with abnormal cell clusters (arrows) and cell-sparse gaps (asterisks). (*F*) HA immunostaining (green) revealed the complete morphology of sparsely-labeled SPNs. Neurons were reconstructed based on confocal Z-stacks. Dendrites are indicated in black. Axons are indicated in blue. In cKO-E, some SPNs were characterized by a dendrite ventrally directed into the intermediate zone (red arrowheads). (*G*) Quantification of SPN morphological subclasses.

To analyze the morphology of single SPNs, we leveraged an *in vivo* genome editing method that targets the actin gene *Actb* for sparse, whole-cell labeling. We used CRISPR-Cas9 to generate a DNA break at the C-terminus of *Actb* and a homology-independent repair template containing a 3xHA epitope tag (54). Non-homology end joining repair that incorporates the template in forward orientation would lead to in-frame expression of ACTB-3xHA. To perform this assay *in vivo*, we used *in utero* electroporation (IUE) to co-transfect CRISPR-Cas9, repair template, and CAG-mTagBFP2 into cortical NPCs at E11.5, at the peak of subplate neuronogenesis (schematized in **Fig. 5*D***). Analyzed at E16.5, mTagBFP2 expression revealed successful targeting of SPNs (**Fig. 5*E***). In control, mTagBPF2-labeled neurons were organized in a continuous narrow band positioned beneath cortical plate. In cKO-E, mTagBPF2-labeled neurons showed aberrant clustering and gaps.

Next, we used anti-HA immunostaining to reveal the morphology of sparsely-labeled SPNs. Reconstruction of confocal z-stacks revealed a diversity of SPN morphologies in control, including pyramidal, horizontal, and multipolar (**Fig. 5 *F*** and ***G***). Diverse subclasses in SPN morphology have been previously documented (7, 65-67), although embryonic SPN morphologies have not been extensively characterized. In cKO-E, morphologies of SPNs were also diverse; however, many deviated from morphological subclasses observed in normal subplate. In addition, some SPNs were characterized by a ventrally-directed dendrite extending into intermediate zone (red arrowheads, **Fig. 5*F***). Notably, unlike SPNs, the morphology of pyramidal neurons in cKO-E was indistinguishable from control (***SI Appendix*, Fig. S5*B***). Thus, our data revealed a cell type-dependent *Arid1a* function in morphogenesis of SPNs.

### Disrupted extracellular matrix and subplate-thalamocortical axon co-fasciculation following *Arid1a* deletion

SPNs are the first neurons to extend corticofugal axons. These descending axons contribute to guidance of thalamocortical ascending axons via co-fasciculation in a model known as the “handshake hypothesis” (26, 68, 69) (schematized in **Fig. 6*A***). To visualize corticofugal axons from SPNs, we used the Tg(*Golli-tau-EGFP*) transgene, which expresses TAU (τ)EGFP in SPNs and L6 neurons (70). In E15.5 control, an abundance of τEGFP+ axons had extended across the PSB. In the subpallium, these descending axons were closely apposed with NTNG1+ thalamocortical axons in a manner consistent with co-fasciculation (solid arrowheads, **Fig. 6*B***). In contrast, in cKO-E, innervation of the subpallium by τEGFP+ axons was attenuated and no co-fasciculation with NTNG1+ axons was found (open arrowheads, **Fig. 6*B***). By E17.5, extensive co-fasciculation of τEGFP+ and NTNG1+ axons was present in control, but was largely absent from cKO-E (**Fig. 6*C***). In addition, in P0 cKO-E, some NTNG1+ axons followed the aberrant trajectories of misrouted τEGFP+ axons, whereas other NTNG1+ axons, without co-fasciculation, were misrouted (***SI Appendix*, Fig. S6*A***). Concomitant with loss of co-fasciculation in cKO-E, NTNG1+ thalamocortical axons were consistently unable to correctly traverse the PSB (arrows, **Fig. 6 *B*** and ***C***, and ***SI Appendix* Fig. S6*B***, n=5/5 animals), entered the cortex via a narrow medial path, became defasciculated in intermediate zone, and prematurely invaded cortical plate. Notably, earlier in development in cKO-E, thalamocortical axons did not show misrouting defects at E13.5, prior to the expected crossing of the PSB at E14.5 (***SI Appendix*, Fig. S6*C***), suggesting that their misrouting followed loss of co-fasciculation with subplate axons. Our data are consistent with the model posited by the “handshake hypothesis” (7).

**Fig. 6.**
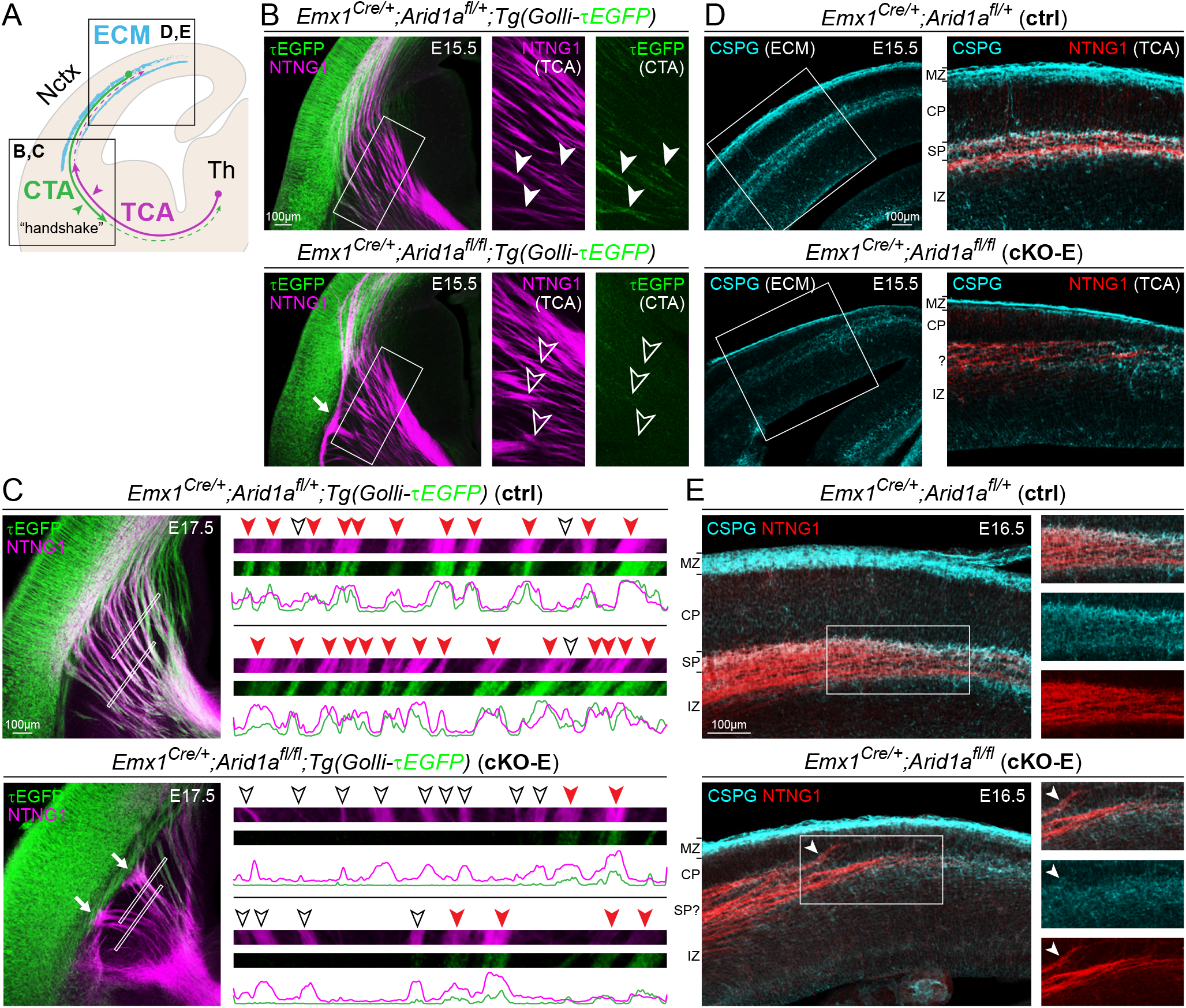
Aberrant subplate neuron axon projection and extracellular matrix following *Arid1a* deletion. (*A*) Schematic illustration of SPN functions. (*B* and *C*) NTNG1 immunostaining (magenta) on E15.5 (*B*) and E17.5 (*C*) ctrl and cKO-E brains carrying the *Golli-τEGFP* transgene. In E15.5 ctrl, τEGFP+ (green) descending axons from SPNs closely co-fasciculated (solid arrowheads) with ascending NTNG1+ thalamocortical axons (magenta). In E15.5 cKO-E, τEGFP+ axons largely have not crossed the pallial-subpallial boundary (PSB). NTNG1+ thalamocortical axons, without co-fasciculation with τEGFP+ axons (open arrowheads), did not cross the PSB along the normal trajectory and formed an aberrant bundle parallel to the boundary (solid arrow) in cKO-E. In E17.5 ctrl, analysis of co-fasciculation (insets) revealed frequent co-fasciculation (red arrowheads) of τEGFP+ and NTNG1+ axons, which is consistent with the “handshake hypothesis”. In E17.5 cKO-E, most NTNG1+ thalamocortical axons did not co-fasciculate with τEGFP+ corticothalamic axons (open arrowheads) and were unable to cross the PSB (solid arrows). (*D* and *E*) CSPG (cyan) and NTNG1 (red) immunostaining on E15.5 (*D*) and E16.5 (*E*) ctrl and cKO-E brain sections. In ctrl, NTNG1+ thalamocortical axons tangentially traversed the embryonic cortex within a subplate/intermediate zone corridor neatly delineated by the extracellular matrix component CSPG. In cKO-E, CSPG expression was reduced and the CSPG corridor had collapsed. NTNG1+ thalamocortical axons were not confined within the subplate/intermediate zone, deviated from their normal trajectory, and prematurely invaded cortical plate (arrowhead). CTA, corticothalamic axon; ECM, extracellular matrix

SPNs are rich in extracellular matrix (71, 72) and secreted molecules derived from SPNs are thought to signal the pathfinding of axons traversing the subplate (73). During normal pathfinding, thalamocortical axons innervate the cortex via a white matter corridor delineated by extracellular matrix component chondroitin sulfate proteoglycan (CSPG) (30), which we observed in E15.5 and E16.5 control (**Fig. 6 *D*** and ***E***). In cKO-E, CSPG expression was reduced and the CSPG corridor had collapsed (**Fig. 6 *D*** and ***E***). Concomitant with CSPG corridor disruption, NTNG1+ thalamocortical axons were not confined within white matter, deviated from their normal trajectory, and invaded cortical plate. Together, our analyses revealed deficits in SPN morphogenesis, subplate axon co-fasciculation with thalamocortical axons, and subplate extracellular matrix following *Arid1a* deletion.

### Subplate-spared cortical plate deletion of *Arid1a* extensively corrected axon misrouting

To empirically test the hypothesis that SPNs mediate the axon guidance functions of *Arid1a*, we sought to determine whether *Arid1a* expression in SPNs was sufficient to support axon pathfinding from cortical neurons that lack *Arid1a*. To do this, we generated an *Arid1a* cKO using Tg(*hGFAP-Cre*) (74). Tg(*hGFAP-Cre*) mediates Cre recombination in cortical NPCs starting at E12.5, after the majority of SPNs have been generated (**Fig. 7*A*** and ***SI Appendix*, Fig. S7*A***); therefore, *Arid1a* would be deleted from cortical plate neurons, whereas SPNs would be spared. Subplate-spared cortical deletion was confirmed by ARID1A immunostaining in these mutants (Tg[*hGFAP-Cre*];*Arid1a*^*fl/fl*^, **cKO-hG**) (**Fig. 7*A***). Importantly, Tg(*hGFAP-Cre*) mediated *Arid1a* deletion from the majority of L6 neurons by E13.5 (**Fig. 7*A*** and ***SI Appendix*, Fig. S7 *B*** and ***C***), thus uncoupling the effects of SPNs from molecularly-similar L6 neurons. In P0 cKO-hG, sparing SPNs from *Arid1a* deletion led to correct anatomical formation of the subplate band (**Fig. 7 *B*** and ***C***). Next, we analyzed cortical axon tracts using Cre-dependent fluorescent reporter *ROSA*^*tdTomato*^ (75) (**Fig. 7*B***). Remarkably, in cKO-hG, axons arising from tdTomato*-*labeled, *Arid1a*-deleted, cortical plate neurons correctly formed the corpus callosum and projected into contralateral cortex (**Fig. 7 *B*** and ***C***). In addition to intracortical axons, thalamocortical axons were also normal in pathfinding in cKO-hG. At P7, CO staining on flattened cortices revealed typical organization of barrel cortex (**Fig. 7*D*** and ***SI Appendix*, Fig. S7*D***, n=3/3 animals). Normal barrel development was confirmed by NTNG1 in cKO-hG (**Fig. 7*D***). Thus, *Arid1a* expression in SPNs was sufficient for normal subplate organization, corpus callosum formation, and thalamocortical axon targeting.

**Fig. 7.**
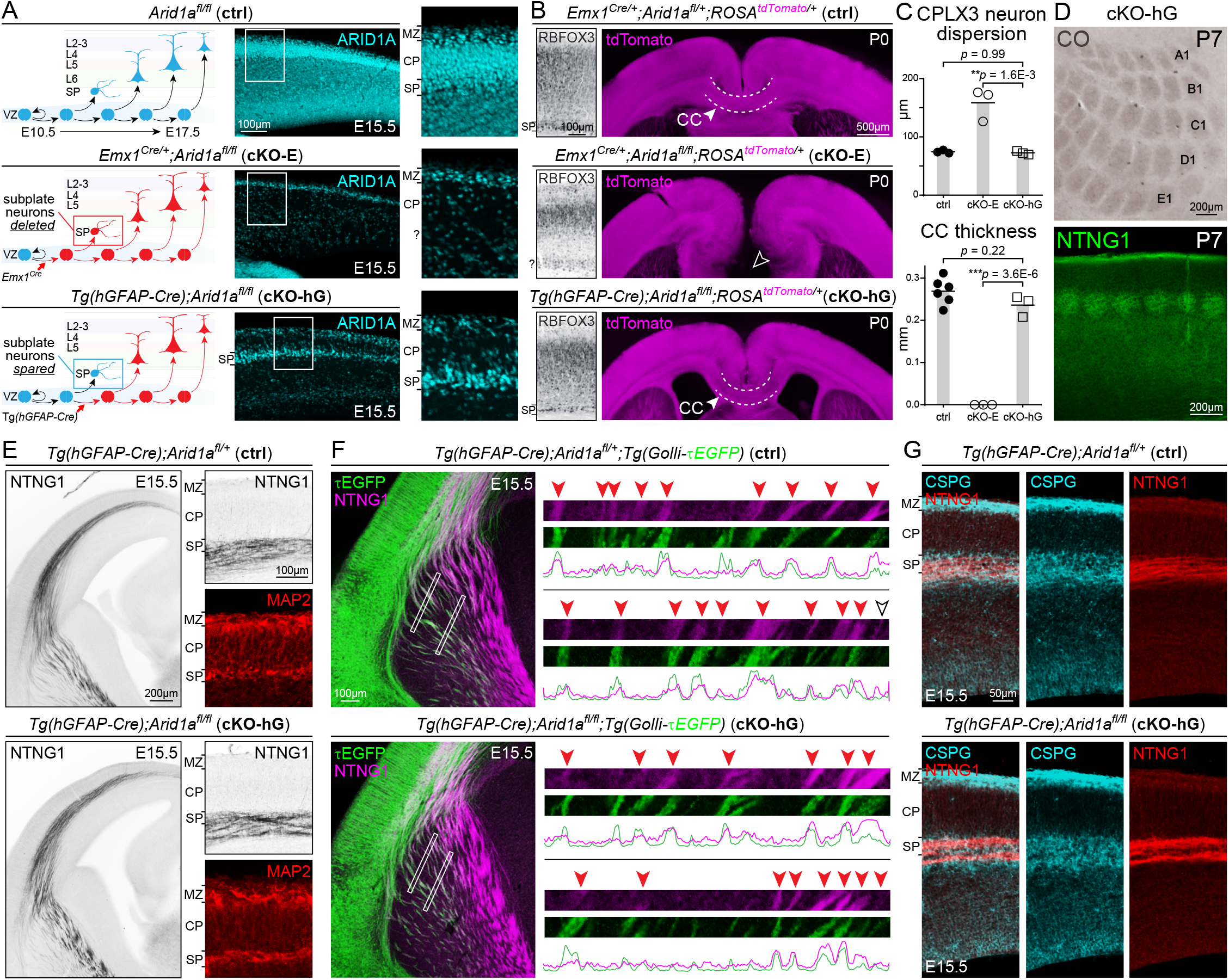
Subplate-spared cortical plate deletion of *Arid1a*. (*A*) Schematic illustration of subplate-spared cortical plate deletion of *Arid1a. Emx1*^*Cre*^ mediates Cre recombination in cortical NPCs starting at E10.5, prior to subplate neuronogenesis. ARID1A immunostaining (cyan) in E15.5 *Emx1*^*Cre/+*^*;Arid1a*^*fl/fl*^ (cKO-E) revealed loss of ARID1A from SPNs and cortical plate neurons. Tg(*hGFAP-Cre*) mediates Cre recombination in cortical NPCs starting at E12.5, after the majority of SPNs have been generated. In E15.5 Tg(*hGFAP-Cre*);*Arid1a*^*fl/fl*^ (cKO-hG), ARID1A was lost from cortical plate neurons, but present in SPNs. (*B*) Subplate and axon tract analyses on P0 ctrl, cKO-E, and cKO-hG brain sections. RBFOX3 immunostaining revealed in cKO-hG an organized, distinct subplate band positioned just beneath the cortical layers that was indistinguishable from ctrl. tdTomato (magenta) was expressed Cre-dependently from *ROSA*^*tdTomato*^, enabling visualization of cortical axons. Agenesis of corpus callosum (open arrowhead) was observed in cKO-E. However, the corpus callosum (CC) was formed without gross defect in cKO-hG (solid arrowhead, n=3/3 animals). (*C*) Quantitative analyses revealed no significant changes in CPLX3+ SPN radial dispersion or corpus callosum thickness at midline in cKO-hG compared to ctrl (data are mean, ANOVA with Tukey’s post hoc test, n≥ 3 animals). (*D*) Whisker barrels in cKO-hG primary somatosensory cortex were visualized by cytochrome oxidase staining (CO, brown) on flattened cortices and NTNG1 immunostaining (green) on coronal sections. Whisker barrels were formed without defect in cKO-hG (n=3/3 animals). (*E*) Analysis of thalamocortical axons and SPNs in E15.5 ctrl and cKO-hG cortex. In cKO-hG, NTNG1+ thalamocortical axons extended along a normal trajectory across the PSB, without forming an aberrant bundle parallel to the boundary. Upon reaching the cortex, NTNG1+ axons were correctly paused within subplate and did not prematurely invade cortical plate in cKO-hG. MAP2 immunostaining (red) revealed in cKO-hG SPNs that were organized within a continuous and clearly delineated layer below cortical plate. (*F*) NTNG1 immunostaining (magenta) on E15.5 ctrl and cKO-hG brains carrying the *Golli-τEGFP* transgene. In cKO-hG, τEGFP+ (green) descending axons from SPNs closely co-fasciculated (red arrowheads) with ascending NTNG1+ thalamocortical axons (magenta). (*G*) CSPG (cyan) and NTNG1 (red) immunostaining on E15.5 ctrl and cKO-hG brain sections. In cKO-hG, NTNG1+ thalamocortical axons travelled within a subplate/intermediate zone corridor neatly delineated by extracellular matrix component CSPG in a manner indistinguishable from ctrl.

Next, we assessed whether correct formation of callosal and thalamocortical tracts in cKO-hG was coincident with normal subplate wiring functions. At E15.5, MAP2 immunostaining in cKO-hG revealed typical organization of SPNs indistinguishable from control (**Fig. 7*E***). NTNG1+ thalamocortical axons entered the cortex along a normal trajectory and without premature invasion of cortical plate (**Fig. 7*E***). *Golli-τEGFP-*labeled corticofugal axons and NTNG1*-*labeled thalamocortical axons showed extensive co-fasciculation in E15.5 cKO-hG (**Fig. 7*F***), which is consistent with normal innervation of the thalamus by descending τEGFP+ axons (***SI Appendix*, Fig. S7*E***). Furthermore, a CSPG corridor properly delineating the path of thalamocortical axons was present in cKO-hG (**Fig. 7*G***). Therefore, despite the absence of ARID1A from cortical plate neurons, subplate expression of ARID1A was sufficient for normal subplate organization, subplate-thalamocortical axon co-fasciculation, and extracellular matrix. Consistent with these wiring functions, subplate *Arid1a* sufficiently enabled normal callosum formation, thalamocortical axon targeting, and whisker barrel development.

Together with our data from *Arid1a* cKO-E, our analysis of *Arid1a* cKO-hG supported examination of cell and non-cell autonomous *Arid1a* functions. In SPNs, *Arid1a* was required for transcription of subplate genes and gave rise to correct SPN organization, co-fasciculation with thalamocortical axons, and extracellular matrix. By regulating the identity and functions of SPNs, *Arid1a* non-cell autonomously controlled the wiring of callosal and thalamocortical connectivities via the axon guidance roles of the subplate (schematized in ***SI Appendix*, Fig. S8**). *Arid1a* is thus a central regulator of multiple subplate-dependent axon guidance mechanisms essential to cortical circuit assembly.

## Discussion

Despite the central role of SPNs in cortical circuit assembly and their potential contribution to neurodevelopmental disorders (5, 23, 25), they are relatively understudied compared to their cortical plate counterparts. Previous studies have focused on subplate-enriched genes (24, 32) and characterized important molecular determinants of SPN specification, migration, and axon projection (33-39). The severe axon misrouting phenotypes of subplate ablation (18-20) however, are not broadly recapitulated in these genetic mutants. Here, we leverage cortical *Arid1a* deletion, which causes axon misrouting defects strikingly reminiscent of subplate ablation, to gain mechanistic insights into the non-cell autonomous wiring functions of SPNs in the assembly of cortical connectivities.

Cortical *Arid1a* deletion and previous experimental subplate ablation (18-21) phenotypically converge on misrouted thalamocortical axons, which prematurely invade the cortical plate and ultimately fail to innervate their L4 targets with correct topology. Several aspects of subplate function may contribute to correct thalamocortical axon pathfinding. First, the “handshake hypothesis” posits that close co-fasciculation between the earliest descending subplate axons and ascending thalamocortical axons is important for guidance of both tracts and formation of reciprocal connectivity (26, 68, 76). Following *Arid1a* deletion in cKO-E, subplate corticofugal axons are markedly reduced, and their co-fasciculation with thalamocortical axons is lost. Consistent with the “handshake hypothesis”, thalamocortical axons, in the absence of co-fasciculation, are impaired in their crossing of the PSB, enter the cortex via a narrow medial path, and become defasciculated and misrouted after entering the cortex. These phenotypes are reminiscent of misrouting defects that follow disrupted subplate function (28, 29) or abnormal shifting of subplate due to piriform cortex expansion (77). Second, subplate is characterized by a rich extracellular matrix (4, 48), which can contribute to axon guidance by interacting with growth cones and supporting guidance signaling (5, 78). During circuit formation, cortical afferent and efferent axons extend along a white matter corridor delineated by matrix component CSPG (30). Following cortical *Arid1a* deletion, CSPG expression is reduced and the corridor collapses. Concomitantly, thalamocortical axons become defasciculated and prematurely invade cortical plate, a phenotype reminiscent of subplate ablation (18). Notably, the sparing of SPNs from *Arid1a* deletion in cKO-hG is sufficient to support both subplate-thalamocortical axon co-fasciculation and the CSPG corridor, and enables correct thalamocortical axon pathfinding. The roles of *Arid1a* in thalamocortical tract formation are therefore centered on SPNs. Interestingly, corticothalamic axons from cortical plate neurons are largely intact following *Arid1a* deletion in cKO-E, despite reduced subplate-thalamic axons. Thus, we do not find an *Arid1a*-dependent pioneering role for subplate axons in guiding corticothalamic axons from subsequent cortical plate neurons.

Unlike the better-known roles of SPNs in thalamocortical axon guidance, subplate contribution to intracortical tract development is less established. Early studies suggest that SPNs pioneer corpus callosum formation by extending the first callosal axons (65, 79-81). Some subsequent studies, however, find this possibility to be unlikely (82, 83). We find that pan-cortical *Arid1a* deletion in cKO-E leads to corpus callosum agenesis and mistargeting of intracortical axons. Sparse *Arid1a* deletion, however, does not autonomously misroute callosal axons, indicating that callosal agenesis is a non-cell autonomous consequence of pan-cortical *Arid1a* deletion. Remarkably, subplate expression of *Arid1a* in cKO-hG is sufficient for normal formation of corpus callosum. Thus, we unequivocally establish that subplate function is essential to callosum development. We note that Tg(*hGFAP-Cre*) is active in indusium griseum and glial wedge (84, 85). Thus, it is unlikely that *Arid1a* expression in these structures could contribute to callosum formation in cKO-hG. Diverse developmental disorders are characterized by agenesis or dysgenesis of corpus callosum (86). Our study highlights a potential contribution of subplate dysfunction to callosal defects in disease.

One barrier to molecular study of subplate function is the lack of specific genetic access to embryonic SPNs during critical stages of circuit wiring. Although several published Cre lines show SPN specificity, the onset of Cre expression occurs too late for study of circuit development (87). Here, we describe a genetic strategy to target SPNs. We find that *Emx1*^*Cre*^ mediates gene deletion from all cortical NPCs, including those that give rise to SPNs, whereas Tg(*hGFAP-Cre*) mediates deletion from NPCs after SPNs have been generated. Importantly, Tg(*hGFAP-Cre*) mediates recombination in a majority of L6 neurons, thereby enabling potential effects of SPNs to be uncoupled from closely-related L6 neurons. By comparing “pan-cortical deletion” (*Emx1*^*Cre*^) versus “subplate-spared deletion” (Tg[*hGFAP-Cre*]), this approach enables interrogation of gene necessity and sufficiency in subplate-mediated circuit wiring.

In cortical NPCs, we find that *Arid1a* deletion leads to a selective disruption in SPN gene expression. Despite ubiquitous ARID1A expression during cortical development, the effects of *Arid1a* deletion are surprisingly cell type-dependent. A recent study showed that *Arid1b* expression following *Arid1a* depletion is sufficient to support BAF complex function (49). In our cKO-E, the expression of *Arid1b* may have attenuated the effects of *Arid1a* loss from cortical plate neurons, thus contributing to subplate-selective deficits.

Recent human genetic findings have convergently implicated altered chromatin function in disorders of brain development (88, 89). These studies have identified loss-of-function mutations in *ARID1A* in intellectual disability, autism spectrum disorder, and Coffin-Siris syndrome, a developmental disorder characterized by callosal dysgenesis (40). The mechanisms by which chromatin dysregulation contribute to brain disorders are an active field of study. An important implication of our work is that deficits in SPNs may be an underappreciated contributor to neural circuit miswiring in brain disorders associated with chromatin dysregulation. In addition, our study highlights the possibility that chromatin regulation can non-cell autonomously contribute to neural circuit development and disorders thereof.

## Materials and Methods

Further experimental details can be found in *SI Appendix, Supplementary Materials and Methods*.

### Mice

All experiments were carried out in compliance with ethical regulations for animal research. Our study protocol was reviewed and approved by the University of Michigan Institutional Animal Care & Use Committee. *Arid1a, Emx1*^*Cre*^, *ROSA*^*mTmG*^, Tg(*hGFAP-Cre*), and *ROSA*^*tdTomato*^ mice were obtained from the Jackson laboratory (Stock nos. 027717, 005628, 007676, 004600, and 007914). Tg(*Lpar1-EGFP*) was purchased from the MMRRC (MMRRC:030171). Tg(*Golli-tau-EGFP*) was a kind gift from Anthony Campagnoni. Date of vaginal plug was considered embryonic day 0.5. Genotyping primers can be found in *SI Appendix*, Table S1.

### Immunostaining and Imaging

Brains were isolated and fixed in 4% PFA overnight at 4° C, embedded in 4% agarose, and vibratome-sectioned at 70 µm. Free-floating sections were incubated with primary antibodies overnight at 4° C, washed 3×5min in PBS, and incubated with corresponding fluorescent secondary antibodies and DAPI for 1h at RT. Images were acquired on an Olympus SZX16, FV1000, or FV3000. Antibodies can be found in *SI Appendix*, Table S2.

### *In Utero* Electroporation

Approximately 2 uL of 1.5 μg/μL CAG-sxiCre-EGFP, or a mix of Actb_NHEJ_3xHA, eSpCas9opt1.1-Actb_gRNA, and CAG-mTagBFP2 each at 1 μg/μL was injected into lateral ventricles. Plasmids were transferred to NPCs in the VZ by electroporation (five 45-ms pulses of 27 V at 950-ms intervals).

### ClickSeq

RNA-seq libraries were generated by Click-Seq (52) from 600 ng of purified neocortical RNA and sequenced at the University of Michigan sequencing core on the Illumina NextSeq550 platform (75 cycle, high output). ddPCR primers and probes can be found in *SI Appendix*, Table S3.

## Acknowledgements

We thank members of the Kwan laboratory for discussions and comments on the study, for critical reading of the manuscript, for scientific discussions, and colleagues in the MNI and Dept. of Human Genetics for insightful suggestions. This work was supported by National Institutes of Health (R01 NS097525 to K.Y.K., F31 NS110206 to D.Z.D., T32 GM007544 to O.H.F.), the Brain Research Foundation (BRFSG-2016-04 to K.Y.K.), March of Dimes Foundation (#5-FY15-33 to K.Y.K.), and Simons Foundation Autism Research Initiative (402213 and 324586 to K.Y.K.).

## Author Contributions

K.Y.K. conceived the study. D.Z.D. and K.Y.K designed experiments. D.Z.D., M.M.L., and A.S. performed experiments. D.Z.D., A.Q., Y.Q., O.H.F., and K.Y.K. analyzed data. D.Z.D. and K.Y.K. wrote the paper.

## Competing Interests

The authors declare no competing financial interests.

## Notes

### Competing Interest Statement

The authors have declared no competing interest.

## References

1. K. L. Allendoerfer, C. J. Shatz, The subplate, a transient neocortical structure: its role in the development of connections between thalamus and cortex. Annu Rev Neurosci 17, 185–218 (1994).

2. S. McConnell, A. Ghosh, C. Shatz, Subplate neurons pioneer the first axon pathway from the cerebral cortex. Science 245, 978–982 (1989).

3. S. McConnell, A. Ghosh, C. Shatz, Subplate pioneers and the formation of descending connections from cerebral cortex. J Neurosci 14, 1892–1907 (1994).

4. I. Kostović, P. Rakic,Developmental history of the transient subplate zone in the visual and somatosensory cortex of the macaque monkey and human brain. J Comp Neurol 297, 441–470 (1990).

5. A. Hoerder-Suabedissen, Z. Molnár, Development, evolution and pathology of neocortical subplate neurons. Nat Rev Neurosci 16, 133–146 (2015).

6. W. Z. Wang et al., Subplate in the developing cortex of mouse and human. J Anat 217, 368–380 (2010).

7. Z. Molnár, R. Adams, A. M. Goffinet, C. Blakemore, The role of the first postmitotic cortical cells in the development of thalamocortical innervation in the reeler mouse. J Neurosci 18, 5746–5765 (1998).

8. Z. Molnár, H. J. Luhmann, P. O. Kanold, Transient cortical circuits match spontaneous and sensory-driven activity during development. Science 370 (2020).

9. I. Kostović, M. E. Molliver, A new interpretation of the laminar development of cerebral cortex: synaptogenesis in different layers of neopallium in the human fetus. Anat Rec 178, 395 (1974).

10. I. Kostović, The enigmatic fetal subplate compartment forms an early tangential cortical nexus and provides the framework for construction of cortical connectivity. Prog Neurobiol 194, 101883 (2020).

11. Z. Molnár et al., New insights into the development of the human cerebral cortex. J Anat 235, 432–451 (2019).

12. K. Y. Kwan, N. Sestan, E. S. Anton, Transcriptional co-regulation of neuronal migration and laminar identity in the neocortex. Development 139, 1535–1546 (2012).

13. P. Rakic, Neurons in rhesus monkey visual cortex: systematic relation between time of origin and eventual disposition. Science 183, 425–427 (1974).

14. P. O. Kanold, H. J. Luhmann, The subplate and early cortical circuits. Annu Rev Neurosci 33, 23–48 (2010).

15. E. Friauf, S. K. McConnell, C. J. Shatz, Functional synaptic circuits in the subplate during fetal and early postnatal development of cat visual cortex. J Neurosci 10, 2601–2613 (1990).

16. J. M. Wess, A. Isaiah, P. V. Watkins, P. O. Kanold, Subplate neurons are the first cortical neurons to respond to sensory stimuli. Proc Natl Acad Sci U S A 114, 12602–12607 (2017).

17. C. Ohtaka-Maruyama et al., Synaptic transmission from subplate neurons controls radial migration of neocortical neurons. Science 360, 313–317 (2018).

18. A. Ghosh, C. J. Shatz, A role for subplate neurons in the patterning of connections from thalamus to neocortex. Development 117, 1031–1047 (1993).

19. A. Ghosh, A. Antonini, S. K. McConnell, C. J. Shatz, Requirement for subplate neurons in the formation of thalamocortical connections. Nature 347, 179–181 (1990).

20. A. Ghosh, C. J. Shatz, Involvement of subplate neurons in the formation of ocular dominance columns. Science 255, 1441–1443 (1992).

21. P. O. Kanold, C. J. Shatz, Subplate neurons regulate maturation of cortical inhibition and outcome of ocular dominance plasticity. Neuron 51, 627–638 (2006).

22. P. O. Kanold, P. Kara, R. C. Reid, C. J. Shatz, Role of subplate neurons in functional maturation of visual cortical columns. Science 301, 521–525 (2003).

23. M. Serati et al., The Role of the Subplate in Schizophrenia and Autism: A Systematic Review. Neuroscience 408, 58–67 (2019).

24. A. Hoerder-Suabedissen et al., Expression profiling of mouse subplate reveals a dynamic gene network and disease association with autism and schizophrenia. Proc Natl Acad Sci U S A 110, 3555–3560 (2013).

25. I. Kostović, M. Judaš, G. Sedmak, Developmental history of the subplate zone, subplate neurons and interstitial white matter neurons: relevance for schizophrenia. Int J Dev Neurosci 29, 193–205 (2011).

26. C. Blakemore, Z. Molnár, Factors involved in the establishment of specific interconnections between thalamus and cerebral cortex. Cold Spring Harb Symp Quant Biol 55, 491–504 (1990).

27. Z. Molnár, C. Blakemore, How do thalamic axons find their way to the cortex? Trends Neurosci 18, 389–397 (1995).

28. D. Magnani, K. Hasenpusch-Theil, T. Theil, Gli3 controls subplate formation and growth of cortical axons. Cereb Cortex 23, 2542–2551 (2013).

29. Y. Chen, D. Magnani, T. Theil, T. Pratt, D. J. Price, Evidence that descending cortical axons are essential for thalamocortical axons to cross the pallial-subpallial boundary in the embryonic forebrain. PLoS One 7, e33105 (2012).

30. A. R. Bicknese, A. M. Sheppard, D. D. O’Leary, A. L. Pearlman, Thalamocortical axons extend along a chondroitin sulfate proteoglycan-enriched pathway coincident with the neocortical subplate and distinct from the efferent path. J Neurosci 14, 3500–3510 (1994).

31. D. J. Price, S. Aslam, L. Tasker, K. Gillies, Fates of the earliest generated cells in the developing murine neocortex. J Comp Neurol 377, 414–422 (1997).

32. F. M. Oeschger et al., Gene expression analysis of the embryonic subplate. Cereb Cortex 22, 1343–1359 (2012).

33. W. Han et al., TBR1 directly represses Fezf2 to control the laminar origin and development of the corticospinal tract. Proc Natl Acad Sci U S A 108, 3041–3046 (2011).

34. R. Hevner et al., Tbr1 regulates differentiation of the preplate and layer 6. Neuron 29, 353–366 (2001).

35. S. Y. X. Tiong et al., Kcnab1 Is Expressed in Subplate Neurons With Unilateral Long-Range Inter-Areal Projections. Front Neuroanat 13, 39 (2019).

36. Y. Arai et al., Evolutionary Gain of Dbx1 Expression Drives Subplate Identity in the Cerebral Cortex. Cell Rep 29, 645-658.e645 (2019).

37. L. Ratie et al., Loss of Dmrt5 Affects the Formation of the Subplate and Early Corticogenesis. Cereb Cortex (2019).

38. K. Y. Kwan et al., SOX5 postmitotically regulates migration, postmigratory differentiation, and projections of subplate and deep-layer neocortical neurons. Proc Natl Acad Sci U S A 105, 16021–16026 (2008).

39. W. L. McKenna et al., Tbr1 and Fezf2 regulate alternate corticofugal neuronal identities during neocortical development. J Neurosci 31, 549–564 (2011).

40. T. Kosho, N. Okamoto, Genotype-phenotype correlation of Coffin-Siris syndrome caused by mutations in SMARCB1, SMARCA4, SMARCE1, and ARID1A. Am J Med Genet C Semin Med Genet 166c, 262–275 (2014).

41. I. Olave, W. Wang, Y. Xue, A. Kuo, G. R. Crabtree, Identification of a polymorphic, neuron-specific chromatin remodeling complex. Genes Dev 16, 2509–2517 (2002).

42. L. Ho, G. R. Crabtree, Chromatin remodelling during development. Nature 463, 474–484 (2010).

43. E. Y. Son, G. R. Crabtree, The role of BAF (mSWI/SNF) complexes in mammalian neural development. Am J Med Genet C Semin Med Genet 166c, 333-349 (2014).

44. X. Gao et al., ES cell pluripotency and germ-layer formation require the SWI/SNF chromatin remodeling component BAF250a. Proc Natl Acad Sci U S A 105, 6656–6661 (2008).

45. J. A. Gorski et al., Cortical excitatory neurons and glia, but not GABAergic neurons, are produced in the Emx1-expressing lineage. J Neurosci 22, 6309–6314 (2002).

46. M. D. Muzumdar, B. Tasic, K. Miyamichi, L. Li, L. Luo, A global double-fluorescent Cre reporter mouse. Genesis 45, 593–605 (2007).

47. I. Kostović, P. Rakic, Development of prestriate visual projections in the monkey and human fetal cerebrum revealed by transient cholinesterase staining. J Neurosci 4, 25–42 (1984).

48. I. Kostović, M. Judas, The development of the subplate and thalamocortical connections in the human foetal brain. Acta Paediatr 99, 1119–1127 (2010).

49. M. Trizzino et al., The Tumor Suppressor ARID1A Controls Global Transcription via Pausing of RNA Polymerase II. Cell Rep 23, 3933–3945 (2018).

50. R. Mathur et al., ARID1A loss impairs enhancer-mediated gene regulation and drives colon cancer in mice. Nat Genet 49, 296–302 (2017).

51. N. Antón-Bolaños, A. Espinosa, G. López-Bendito, Developmental interactions between thalamus and cortex: a true love reciprocal story. Curr Opin Neurobiol 52, 33–41 (2018).

52. A. Routh, S. R. Head, P. Ordoukhanian, J. E. Johnson, ClickSeq: Fragmentation-Free Next-Generation Sequencing via Click Ligation of Adaptors to Stochastically Terminated 3’-Azido cDNAs. J Mol Biol 427, 2610–2616 (2015).

53. L. Shi, A. Qalieh, M. M. Lam, J. M. Keil, K. Y. Kwan, Robust elimination of genome-damaged cells safeguards against brain somatic aneuploidy following Knl1 deletion. Nat Commun 10, 2588 (2019).

54. J. M. Keil et al., Symmetric neural progenitor divisions require chromatin-mediated homologous recombination DNA repair by Ino80. Nat Commun 11, 3839 (2020).

55. M. D. Robinson, D. J. McCarthy, G. K. Smyth, edgeR: a Bioconductor package for differential expression analysis of digital gene expression data. Bioinformatics 26, 139–140 (2010).

56. S. A. Yuzwa et al., Developmental Emergence of Adult Neural Stem Cells as Revealed by Single-Cell Transcriptional Profiling. Cell Rep 21, 3970–3986 (2017).

57. L. Loo et al., Single-cell transcriptomic analysis of mouse neocortical development. Nat Commun 10, 134 (2019).

58. S. M. Sunkin et al., Allen Brain Atlas: an integrated spatio-temporal portal for exploring the central nervous system. Nucleic Acids Res 41, D996–d1008 (2013).

59. M. V. Kuleshov et al., Enrichr: a comprehensive gene set enrichment analysis web server 2016 update. Nucleic Acids Res 44, W90–97 (2016).

60. B. J. Molyneaux et al., DeCoN: genome-wide analysis of in vivo transcriptional dynamics during pyramidal neuron fate selection in neocortex. Neuron 85, 275–288 (2015).

61. A. Visel, C. Thaller, G. Eichele, GenePaint.org: an atlas of gene expression patterns in the mouse embryo. Nucleic Acids Res 32, D552–556 (2004).

62. S. Gong et al., A gene expression atlas of the central nervous system based on bacterial artificial chromosomes. Nature 425, 917–925 (2003).

63. A. Hoerder-Suabedissen et al., Novel markers reveal subpopulations of subplate neurons in the murine cerebral cortex. Cereb Cortex 19, 1738–1750 (2009).

64. A. Hoerder-Suabedissen, Z. Molnár, Molecular diversity of early-born subplate neurons. Cereb Cortex 23, 1473–1483 (2013).

65. A. Hoerder-Suabedissen, Z. Molnár, Morphology of mouse subplate cells with identified projection targets changes with age. J Comp Neurol 520, 174–185 (2012).

66. Z. Molnár, R. Adams, C. Blakemore, Mechanisms underlying the early establishment of thalamocortical connections in the rat. J Neurosci 18, 5723–5745 (1998).

67. J. A. De Carlos, D.D. O’Leary, Growth and targeting of subplate axons and establishment of major cortical pathways. J Neurosci 12, 1194–1211 (1992).

68. Z. Molnár, S. Garel, G. López-Bendito, P. Maness, D. J. Price, Mechanisms controlling the guidance of thalamocortical axons through the embryonic forebrain. Eur J Neurosci 35, 1573–1585 (2012).

69. Z. Molnár, C. Blakemore, Lack of regional specificity for connections formed between thalamus and cortex in coculture. Nature 351, 475–477 (1991).

70. M. C. Piñon, A. Jethwa, E. Jacobs, A. Campagnoni, Z. Molnár, Dynamic integration of subplate neurons into the cortical barrel field circuitry during postnatal development in the Golli-tau-eGFP (GTE) mouse. J Physiol 587, 1903–1915 (2009).

71. I. Kostović, I. Išasegi, Ž. Krsnik, Sublaminar organization of the human subplate: developmental changes in the distribution of neurons, glia, growing axons and extracellular matrix. J Anat 235, 481–506 (2019).

72. M. Judas,N.J. Milosević, M. R. Rasin, M. Heffer-Lauc, I. Kostović, Complex patterns and simple architects: molecular guidance cues for developing axonal pathways in the telencephalon. Prog Mol Subcell Biol 32, 1–32 (2003).

73. S. Kondo, H. Al-Hasani, A. Hoerder-Suabedissen, W. Z. Wang, Z. Molnár, Secretory function in subplate neurons during cortical development. Front Neurosci 9, 100 (2015).

74. L. Zhuo et al., hGFAP-cre transgenic mice for manipulation of glial and neuronal function in vivo. Genesis 31, 85–94 (2001).

75. L. Madisen et al., A robust and high-throughput Cre reporting and characterization system for the whole mouse brain. Nat Neurosci 13, 133–140 (2010).

76. R. F. Hevner, E. Miyashita-Lin, J. L. Rubenstein, Cortical and thalamic axon pathfinding defects in Tbr1, Gbx2, and Pax6 mutant mice: evidence that cortical and thalamic axons interact and guide each other. J Comp Neurol 447, 8–17 (2002).

77. A. Em et al., Expansion of the piriform cortex contributes to corticothalamic pathfinding defects in Gli3 conditional mutants. Cerebral cortex (New York, N.Y.: 1991) 25 (2015).

78. M. Tessier-Lavigne, M. Placzek, A. G. Lumsden, J. Dodd, T. M. Jessell, Chemotropic guidance of developing axons in the mammalian central nervous system. Nature 336, 775–778 (1988).

79. J. J. Chun, M. J. Nakamura, C. J. Shatz, Transient cells of the developing mammalian telencephalon are peptide-immunoreactive neurons. Nature 325, 617–620 (1987).

80. A. Antonini, C. J. Shatz, Relation Between Putative Transmitter Phenotypes and Connectivity of Subplate Neurons During Cerebral Cortical Development. Eur J Neurosci 2, 744–761 (1990).

81. L. C. deAzevedo, C. Hedin-Pereira, R. Lent, Callosal neurons in the cingulate cortical plate and subplate of human fetuses. J Comp Neurol 386, 60–70 (1997).

82. H. S. Ozaki, D. Wahlsten, Timing and origin of the first cortical axons to project through the corpus callosum and the subsequent emergence of callosal projection cells in mouse. J Comp Neurol 400, 197–206 (1998).

83. S. E. Koester, D.D. O’Leary, Axons of early generated neurons in cingulate cortex pioneer the corpus callosum. J Neurosci 14, 6608–6620 (1994).

84. C. Benadiba et al., The ciliogenic transcription factor RFX3 regulates early midline distribution of guidepost neurons required for corpus callosum development. PLoS Genet 8, e1002606 (2012).

85. K. M. Smith et al., Midline radial glia translocation and corpus callosum formation require FGF signaling. Nat Neurosci 9, 787–797 (2006).

86. L. K. Paul et al., Agenesis of the corpus callosum: genetic, developmental and functional aspects of connectivity. Nat Rev Neurosci 8, 287–299 (2007).

87. A. Hoerder-Suabedissen et al., Subset of Cortical Layer 6b Neurons Selectively Innervates Higher Order Thalamic Nuclei in Mice. Cereb Cortex 28, 1882–1897 (2018).

88. S. J. Sanders et al., Insights into Autism Spectrum Disorder Genomic Architecture and Biology from 71 Risk Loci. Neuron 87, 1215–1233 (2015).

89. S. De Rubeis et al., Synaptic, transcriptional and chromatin genes disrupted in autism. Nature 515, 209–215 (2014).

