## Supplemental Materials for "Chromatin remodeler *Arid1a* regulates subplate neuron identity and wiring of cortical connectivity"

**This PDF file includes:**

Supplementary Materials and Methods

Figures S1 to S8

Tables S1 to S3

SI References

**Supplementary Materials and Methods**

**Immunostaining and Imaging.** Free-floating sections were blocked and immunostained in blocking solution containing 5% donkey serum, 1% BSA, 0.1% glycine, 0.1% lysine, and 0.3% Triton X-100 (Triton X-100 was excluded from blocking solution when immunostaining for CSPG). Sections were incubated with primary antibodies in blocking solution overnight at 4° C and with secondary antibodies for 1h at RT. Following secondary antibody staining, sections were mounted with VECTASHIELD Antifade Mounting Medium (Vector Laboratories). Images were acquired using an Olympus SZX16 dissecting scope with Olympus U-HGLGPS fluorescent source and Q-Capture Pro 7 software to operate a Q-imaging Regia 6000 camera, an Olympus Fluoview FV1000 confocal microscope with FV10-ASW software, or an Olympus Fluoview FV3000 confocal microscope with FV31S-SW software. Images were processed and quantified in ImageJ and Adobe Photoshop. Primary and secondary antibodies are listed in *SI Appendix,* Table S2.

**Cytochrome Oxidase.** Brains from P7 mutant and control mice were fixed for 2 h at RT in 4% PFA, and the cortices were dissected off, flattened between two glass slides, and fixed overnight at 4° C. Following fixation, flattened cortices were sectioned on a vibratome at 150 µm. Sections were incubated at 37° C overnight in a solution containing 4 g sucrose, 50 mg DAB (Sigma-Aldrich), 15 mg cytochrome C (Sigma-Aldrich) per 100 mL of PBS. Sections were washed with PBS, and imaged with an Olympus SZX16 dissecting scope.

**Statistical Analysis.** Statistical analyses were performed in GraphPad Prism 8 (GraphPad Software). Values were compared using a two-tailed, unpaired Student's *t* test, ANOVA with Tukey's post hoc test, or hypergeometric test with Bonferroni correction. An $\alpha$ of 0.05 was used to determine statistical significance unless otherwise indicated.

**Plasmid Constructs.** For CAG-sxiCre-EGFP, an iCre sequence followed by polyA and flanked by loxP sites was placed immediately downstream of a CAG promoter. An EGFP sequence, WPRE, and polyA were inserted following the iCre/loxP cassette so that Cre expression will lead to self-excision and EGFP expression. CAG-mTagBFP2 was generated by subcloning mTagBFP2 from mTagBFP2-Lifeact-7 (1) (a gift from Michael Davidson, Addgene plasmid #54602) into pCAGEN (2) (a gift from Connie Cepko, Addgene plasmid #11160). To make eSpCas9opt1.1-Actb_gRNA (3), a gRNA sequence targeting the c-terminus of the *Actb* coding sequence (5′-AGTCCGCCTAGAAGCACTTG) was cloned into eSpCas9Opt1.1, which expresses enhanced-specificity Cas9 and an optimized gRNA scaffold. To construct Actb_NHEJ_3xHA (3), tandem HA tag sequences were cloned between 2 *Actb* gRNA recognition sequences, to enable 3xHA insertion into the *Actb* gene following cutting of the endogenous gene and the plasmid by Cas9.

**RNA Purification and Droplet Digital PCR (ddPCR).** Neocortical tissue was microdissected from E15.5 mice. RNA was isolated by resuspending the tissue in TRIzol, homogenizing with metal beads in a bullet blender, isolating the aqueous phase following addition of chloroform and centrifugation for 15 min at >20,000g at 4°C, and eluting purified RNA with DNase/RNase-free water from a Zymo Research Zymo-Spin IC column. Purified RNA was quantified using a Qubit fluorometer. 1 µg of RNA was reverse transcribed using SuperScript II (Invitrogen). Probe sets (PrimeTime qPCR Probe Assays) from Integrated DNA Technologies were utilized with diluted cDNA and the QX200 Droplet Digital PCR System (Bio-Rad) for gene expression level analyses. The ddPCR reaction mixture contained template cDNA, 2xddPCR Supermix (No dUTP) (Bio-Rad), target probes with HEX fluorescence, and control probe against the house-keeping gene *Srp72* with FAM fluorescence. Droplets were generated with the QX200 droplet generator (Bio-Rad), thermocycled, and analyzed using the QX200 droplet reader (Bio-Rad). Primers and probes for ddPCR can be found in *SI Appendix,* Table S3.

**ClickSeq.** Libraries were generated from 600 ng of purified neocortical RNA. Ribosomal RNA was removed from total RNA using NEBNext rRNA Depletion Kit (NEB). ERCC RNA spike-in was included for library quality assessment (Thermo Fisher). SuperScript II (Invitrogen) was used for reverse transcription with 1:30 5 mM

AzdNTP:dNTP and 3′ Genomic Adapter-6N RT primer (GTGACTGGAGTTCAGACGTGTGCTCTTCCGATCTNNNNNN). RNaseH treatment was used to remove RNA template and DNA was purified with DNA Clean and Concentrator Kit (Zymo Research). Azido-terminated cDNA was combined with the click adaptor oligo (/5Hexynyl/NNNNNNNNAGATCGGAAGAGCGTCGTGTAGGGAAAGAGTGTAGATCTCGGTGGTCGCCGTATCATT) and click reaction was catalyzed by addition of ascorbic acid and $Cu^{2+}$, with subsequent purification with DNA Clean and Concentrator Kit. Library amplification was performed using Illumina universal primer (AATGATACGGCGACCACCGAG), Illumina indexing primer (CAAGCAGAAGACGGCATACGAGATNNNNNNGTGACTGGAGTTCAGACGTGT) and the manufacturer's protocols from the 2× One Taq Hot Start Mastermix (NEB). To enrich for amplification products larger than 200 bp, PCR products were purified using Ampure XP (Beckman) magnetic beads at 1.25× ratio. Libraries were analyzed on TapeStation (Agilent) for appropriate quality and distribution and were sequenced at the University of Michigan sequencing core on the Illumina NextSeq 550 platform (75 cycle, high output).

**RNA Sequencing Data Analysis.** RNA-seq data were subject to quality-control check using FastQC v0.11.5 (https://www.bioinformatics.babraham.ac.uk/projects/download.html#fastqc). Adapters were trimmed using cutadapt version 1.13 (http://cutadapt.readthedocs.io/en/stable/guide.html). Processed reads were aligned to GENCODE GRCm38/mm10 reference genome (https://www.gencodegenes.org/mouse/) with STAR(4) (v2.5.2a) and deduplicated according to UMI using UMI-tools (5) (v0.5.3). Read counts were obtained with htseq-count (v0.6.1p1) with intersection-nonempty mode (6). Differential expression was determined with edgeR (7). The *P* value was calculated with likelihood ratio tests and the adjusted *P* value for multiple tested was calculated using the Benjamini-Hochberg procedure, which controls false discovery rate (FDR). Sequencing results were confirmed for 6 genes and normalized to the housekeeping gene *Srp72* using droplet digital PCR with primers and probes in *SI Appendix*, Table S3.

**Single Cell RNA-seq Intersectional Analysis.** Single cell RNA-seq data were obtained (8, 9). To ensure consistency between methods, unclustered data were clustered using standard t-SNE cell-based clustering (10). Raw counts were normalized with K-nearest neighbor smoothing and a Freeman-Tukey transform to improve the signal-to-noise ratio (11). Pearson product-moment correlation coefficients for gene expression were calculated using NumPy (12) and plotted using seaborn (13).

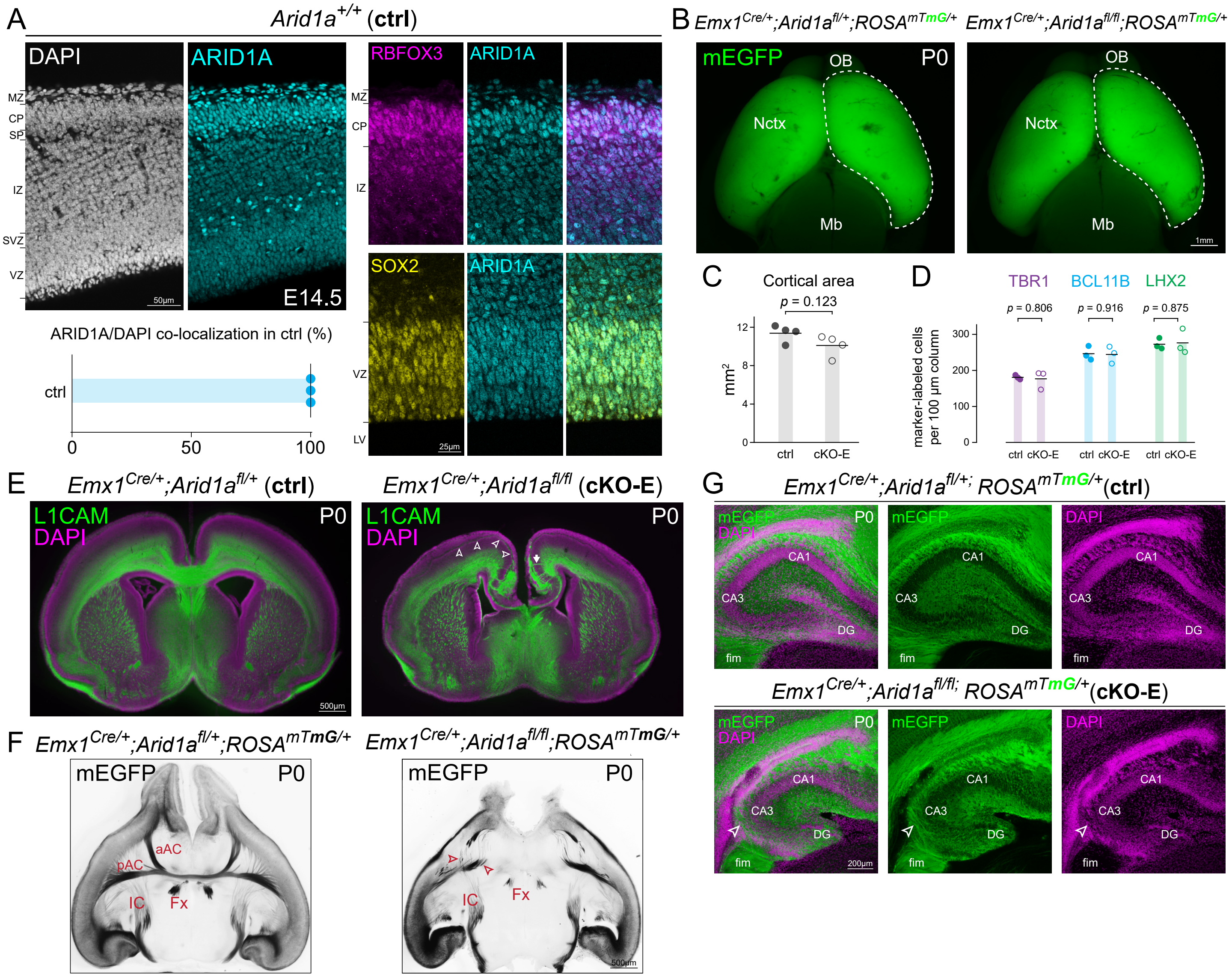


**Fig. S1.** Tract-dependent misrouting of cortical axons following conditional *Arid1a* deletion. (*A*) E14.5 immunostaining of ARID1A (cyan) in control brains revealed ARID1A expression in SOX2+ NPCs (yellow) and RBFOX3+ (NEUN+) neurons (magenta), and complete ARID1A colocalization with DAPI (white) (n=3 animals), supporting ubiquitous expression of ARID1A during cortical development. (*B*) Dorsal view of whole mount P0 ctrl and cKO-E brains. Membrane EGFP (mEGFP, green) was expressed Cre-dependently from *ROSA*$^{mTmG}$. (*C*) Quantitative analysis of cortical area in P0 cKO-E and ctrl (data are mean, two-tailed unpaired *t* test, n=4 animals). (*D*) Quantification of layer marker immunostaining for TBR1 (L6, magenta), BCL11B (L5, cyan), and LHX2 (L2-5, green) revealed no significant changes in P0 cKO-E compared to ctrl (data are mean, two-tailed unpaired *t* test, n=3 animals). (*E*) L1CAM immunostaining (green) of P0 coronal sections confirmed widespread axonal misrouting in cKO-E, including tangentially-directed axons across the upper layers (arrowheads) and radially-directed axons toward the pia (arrow). (*F*) Horizontal sections of P0 ctrl and cKO-E brains. Visualization of cortical projections by *ROSA*$^{mTmG}$ (black) uncovered in cKO-E misrouting of presumptive anterior commissure axons (arrowheads), which failed to cross the midline. Corticofugal axons, however, innervated the internal capsule (IC) in cKO-E without apparent deficit. Hippocampal axons innervating the fornix (Fx) were present in cKO-E, although that innervation was qualitatively reduced compared to ctrl. (*G*) Coronal sections of P0 ctrl and cKO-E hippocampus. DAPI (magenta) and mEGFP (green) revealed hippocampal hypoplasia and axon misrouting (open arrowhead) in cKO-E. CP, cortical plate; LV, lateral ventricle; IZ, intermediate zone; MZ, marginal zone; SP, subplate; SVZ, subventricular zone; VZ, ventricular zone; Mb, midbrain; Nctx, neocortex; OB, olfactory bulb; aAC, anterior branch of the anterior commissure; pAC, posterior branch of the anterior commissure; DG, dentate gyrus; fim, fimbria

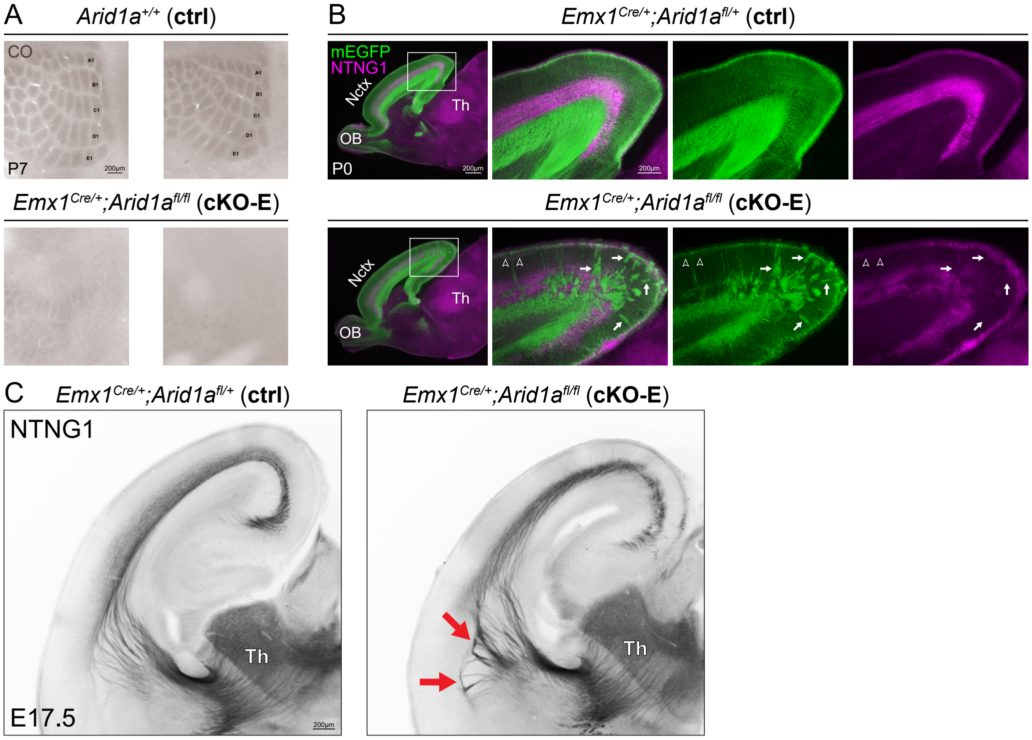


**Fig. S2.** Non-cell autonomous disruption of thalamocortical axon pathfinding following *Arid1a* deletion. (*A*) Cytochrome oxidase (CO, brown) staining of flattened P7 control and cKO-E cortex. Whisker barrels were stereotypically organized in ctrl but largely absent from cKO-E. (*B*) mEGFP (green) and NTNG1 (magenta) immunostaining on sagittal sections of P0 ctrl and cKO-E brains. Both mEGFP+ cortical axons and NTNG1+ thalamocortical axons contributed to the tangentially-directed aberrant axons in the upper layers (open arrowheads) in cKO-E. Only mEGFP+ cortical axons were misrouted into radially-directed bundles toward the pia (arrows). (*C*) NTNG1 immunostaining (black) on coronal sections of E17.5 ctrl and cKO-E brains. In cKO-E, NTNG1+ thalamocortical axons failed to correctly cross the pallial-subpallial boundary (PSB), formed dense axon bundles parallel to the PSB (arrows), and entered the cortex via an aberrant medial path. Th, thalamus

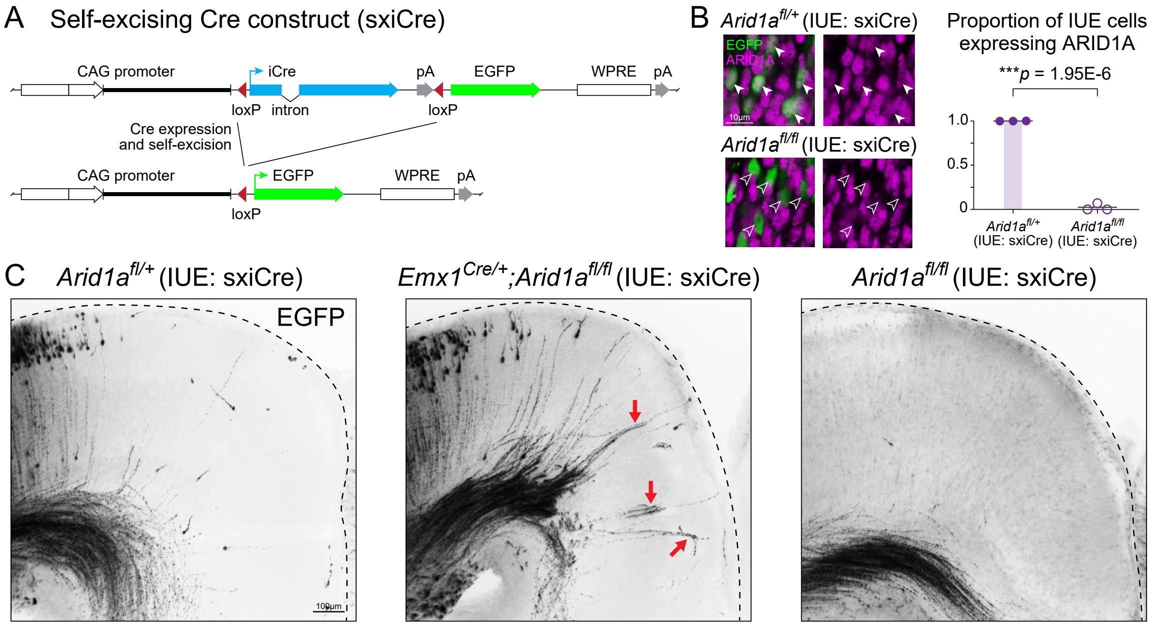


**Fig. S3.** Correct callosal axon targeting following sparse deletion of *Arid1a*. (*A*) Schematic illustration of a self-excising Cre expression EGFP reporter construct (sxiCre) used for sparse deletion of *Arid1a* via *in utero* electroporation (IUE). (*B*) Successful deletion of *Arid1a* in *Arid1a*$^{fl/fl}$ mice (without genetic Cre) following sxiCre IUE was confirmed by ARID1A (magenta) and EGFP (green) immunostaining. ARID1A was lost (open arrowheads) from 97.67% of EGFP+ transfected cells in *Arid1a*$^{fl/fl}$ mice (data are mean, two-tailed unpaired *t* test, n=3 animals). (*C*) EGFP immunostaining (black) on coronal sections of P0 brains after IUE. Following broad genetic *Arid1a* deletion in cKO-E, but not sparse *Arid1a* deletion by sxiCre IUE, transfected neurons extended aberrant radially-directed axons toward the pia (arrows) reminiscent of *ROSA*$^{mTmG}$ axon analysis in cKO-E.

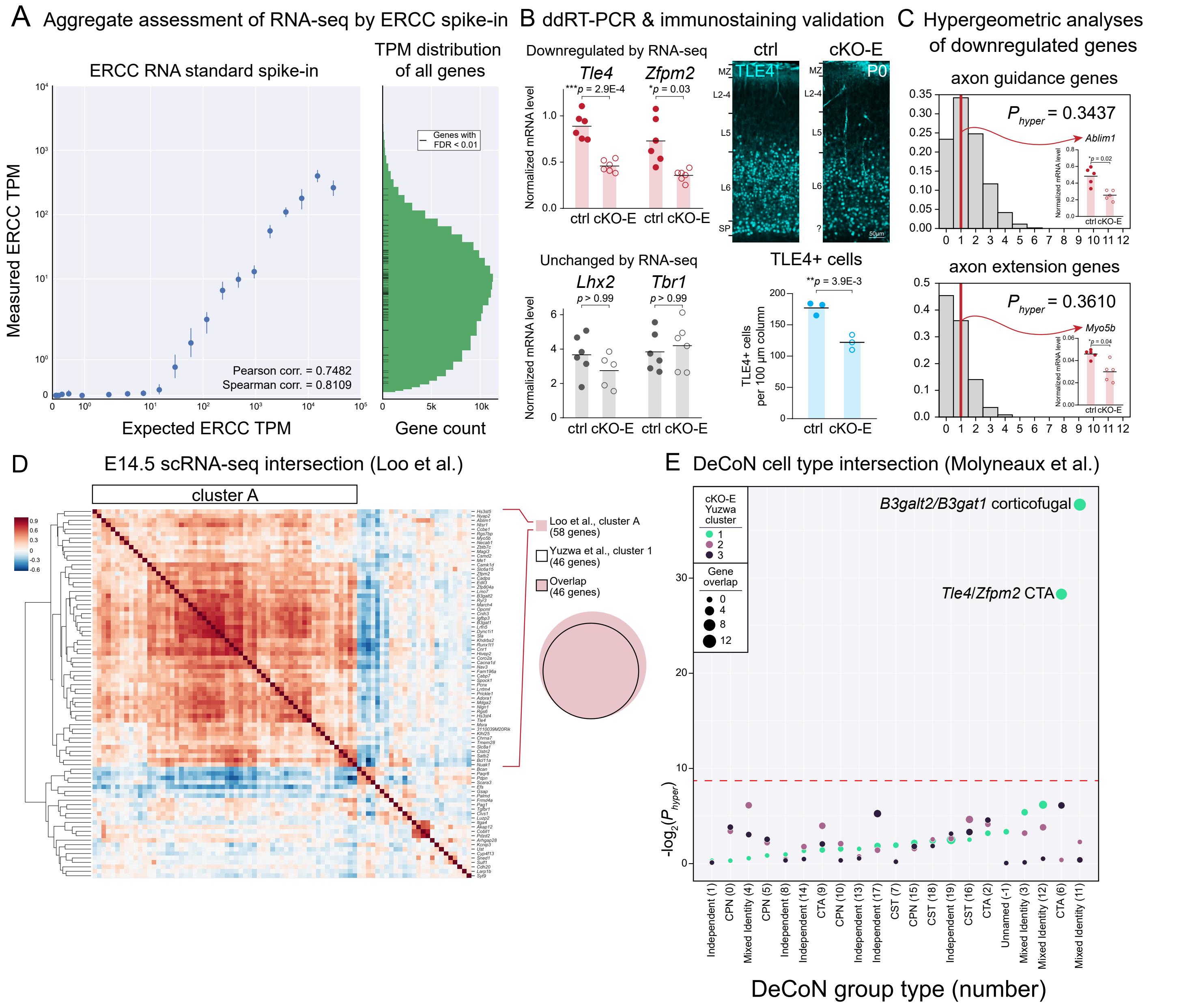


**Fig. S4.** Selective disruption of subplate neuron gene expression following *Arid1a* deletion. (*A*) Aggregate assessment of ERCC spike-in standards in UMI RNA-seq revealed excellent quantification over a broad range of expression levels. All differentially expressed genes were within the dynamic range of UMI RNA-seq. (*B*) Digital droplet (dd)RT-PCR validation of two genes that were downregulated in cKO-E based on RNA-seq (*Tle4, Zfpm2*) and two genes unchanged in cKO-E based on RNA-seq (*Lhx2, Tbr1*) (data are mean, two-tailed *t* test with Bonferroni correction, *Tle4*, *Zfpm2*, and *Tbr1*: n=6 animals, *Lhx2*: ctrl n=6, cKO-E n=5 animals). Immunostaining for TLE4 (cyan) in P0 ctrl and cKO-E cortex confirmed a significant loss of TLE4+ cells in cKO-E (data are mean, two-tailed *t* test, n=3 animals). (*C*) Hypergeometric analysis of genes significantly downregulated in cKO-E (FDR < 0.01) revealed no significant overrepresentation of axon guidance (GO:0007411) or axon extension (GO:0048675) genes (hypergeometric test, Bonferroni correction, $\alpha$ = 0.025). *Ablim1* (axon guidance) and *Myo5b* (axon extension) downregulation in E15.5 cKO-E cortex was confirmed by ddRT-PCR (data are mean, two-tailed *t* test with Bonferroni correction, n=5 animals). (*D*) Intersectional analysis of cKO-E downregulated genes (FDR < 0.01) with single-cell RNA-seq from wildtype E14.5 cortex (Loo et al., 2019) (9) revealed a cluster of 58 genes highly co-expressed in single cells (cluster A). Cluster A encompassed all 46 genes from cluster 1 (Yuzwa et al., 2017 scRNA-seq) (8), providing orthogonal support that gene expression was preferentially affected within a specific cell type in cKO-E. (*E*) Intersection of cKO-E downregulated genes in cluster 1 (Yuzwa et al., 2017 scRNA-seq) (8) with cell type-specific RNA-seq data (Molyneaux et al., 2015) (14) revealed overrepresentation of corticothalamic group 6 and corticofugal group 11, which likely comprise subplate neurons based on marker membership, but no overrepresentation of other cell types (e.g. CPN, CST). Intersectional analyses of cluster 2 and 3 genes revealed no overrepresentation of any cell type (hypergeometric test, Bonferroni correction, $\alpha$ = 0.00238). TPM, transcripts per million

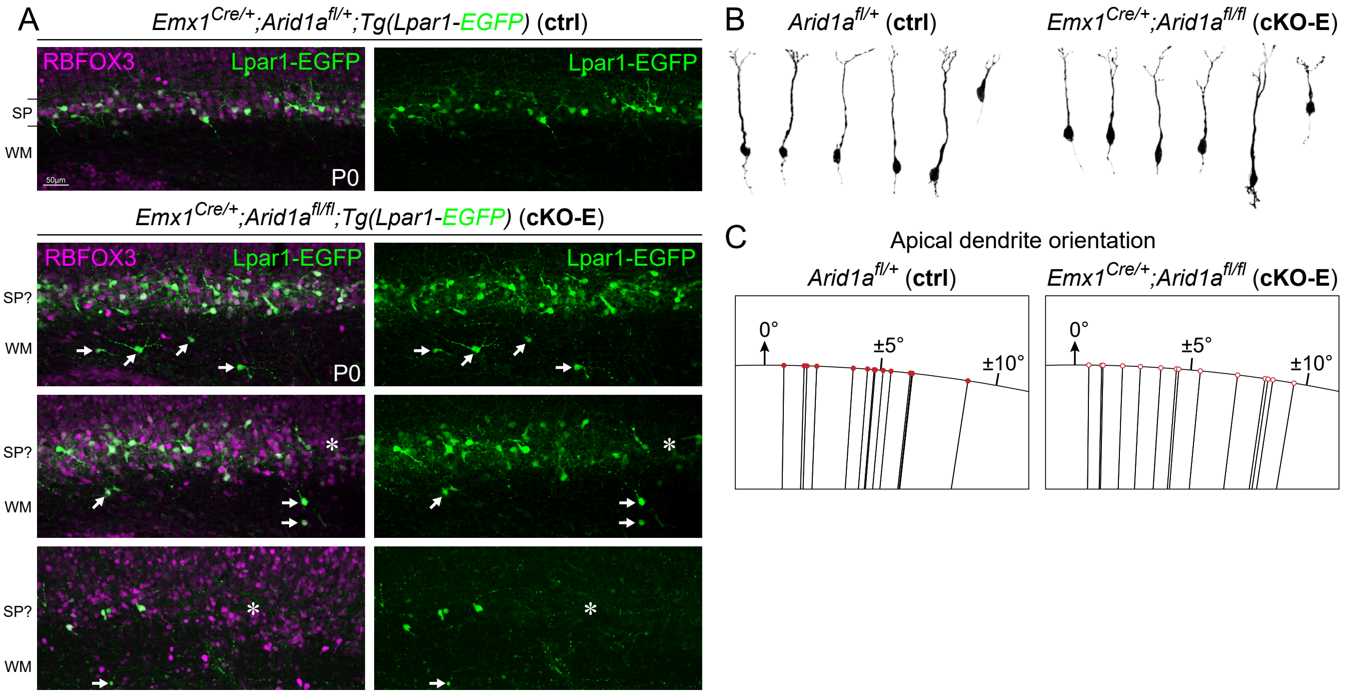


**Fig. S5.** Disrupted subplate organization and subplate neuron morphology following *Arid1a* deletion. (*A*) RBFOX3 (NEUN) immunostaining on P0 ctrl and cKO-E brains carrying the *Lpar1-EGFP* transgene. In cKO-E, RBFOX3+ (magenta) and Lpar1-EGFP+ (green) neurons were not tightly organized into a discreet, continuous subplate band beneath the cortical plate. Lpar1-EGFP+ neurons were aberrantly positioned in white matter (WM, arrows), and the Lpar1-EGFP+ subplate band was characterized by cell-sparse gaps (asterisk). (*B*) Morphological analysis of ACTB-3xHA-tagged pyramidal neurons at E16.5. No robust morphological differences were found between cKO-E and ctrl pyramidal neurons. (*C*) Quantitative analysis of pyramidal neuron apical dendrite orientation at E16.5. No significant difference in apical dendrite orientation was found between cKO-E and ctrl ($p$ = 0.74, two-tailed unpaired $t$ test, n=14 neurons).

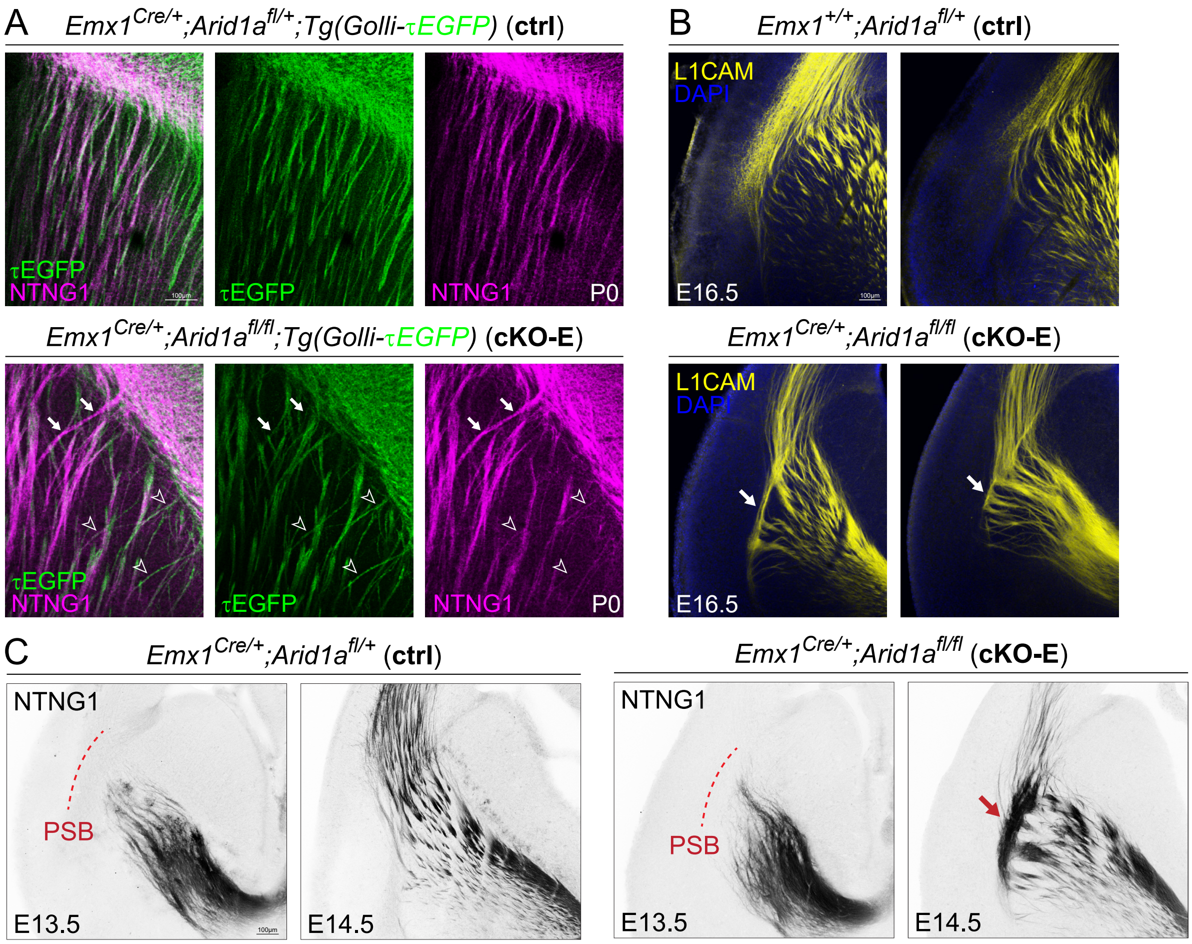


**Fig. S6.** Aberrant subplate neuron axon projection and extracellular matrix following *Arid1a* deletion. (*A*) NTNG1 immunostaining (magenta) on horizontal sections of P0 ctrl and cKO-E brains carrying the *Golli-τEGFP* transgene. In cKO-E, some NTNG1+ thalamocortical axons followed the aberrant trajectories of misrouted τEGFP+ (green) corticofugal axons (open arrowheads). Some NTNG1+ axon bundles, in the absence of co-fasciculation with τEGFP+ axons, were misrouted (solid arrows). (*B*) L1CAM immunostaining (yellow) on coronal E16.5 sections revealed in cKO-E consistent misrouting of axons into aberrant bundles that ran parallel to the pallial-subpallial boundary (PSB, arrows). (*C*) NTNG1 immunostaining (black) on coronal sections of ctrl and cKO-E. At E13.5, NTNG1+ thalamocortical axons had not crossed the PSB in ctrl. In cKO-E, thalamocortical axons had not shown any misrouting deficits. At E14.5, thalamocortical axons abundantly crossed the PSB in ctrl. In cKO-E, thalamocortical axons had started to show characteristic misrouting and defective crossing of the PSB (arrow).

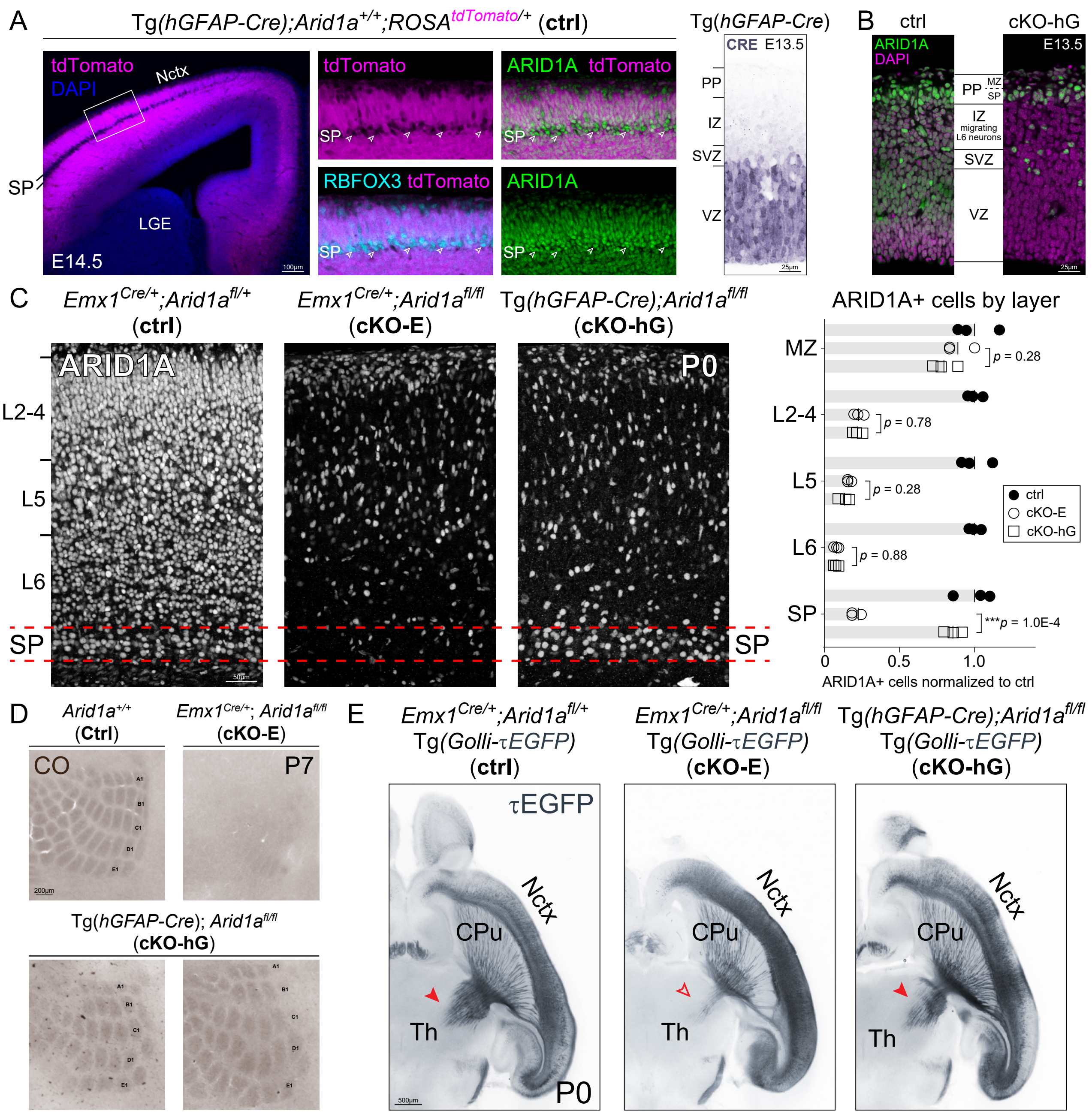


**Fig. S7.** Subplate-spared cortical plate deletion of *Arid1a*. (*A*) Analysis of Tg(*hGFAP-Cre*)-mediated recombination. In E14.5 Tg(*hGFAP-Cre*), analysis using the Cre-dependent reporter *ROSA*$^{tdTomato}$ revealed an absence of tdTomato (magenta) from the subplate (SP) band. Co-immunostaining revealed an absence of tdTomato from RBFOX3+ (cyan) subplate neurons (arrowheads, inset), which expressed ARID1A (green). At E13.5, CRE (blue) was abundant in VZ NPCs, but not in postmitotic neurons in IZ or PP. (*B*) At E13.5, ARID1A (green) was widely expressed in ctrl cortex. In cKO-hG, ARID1A was present in the PP, where future subplate neurons were located, consistent with a sparing of subplate neurons by Tg(*hGFAP-Cre*). In contrast, ARID1A was absent from migrating L6 neurons in IZ and from NPCs in VZ. (*C*) ARID1A immunostaining (white) on coronal sections of P0 ctrl, cKO-E, and cKO-hG brains. ARID1A was present in subplate (red lines) in ctrl and absent from subplate in cKO-E. In cKO-hG, ARID1A expression was present in subplate, confirming sparing of subplate neurons from *Arid1a* deletion. The loss of ARID1A expression from L2-6 neurons, however, was not significantly different between cKO-E and cKO-hG (data are mean, two-tailed unpaired *t* test, n=3 animals). (*D*) Cytochrome oxidase (brown) staining on flattened P7 ctrl, cKO-E, and cKO-hG cortices. Whisker barrel organization was lost from cKO-E, but was unaffected in cKO-hG. (*E*) Horizontal sections of P0 ctrl, cKO-E, and cKO-hG brains carrying the *Golli-τEGFP* transgene. Compared to ctrl, innervation of thalamus (Th) by τEGFP+ axons were qualitatively reduced in cKO-E (open arrowhead), but not in cKO-hG (solid arrowhead). LGE, lateral ganglionic eminence; PP, preplate; CPu, caudate putamen

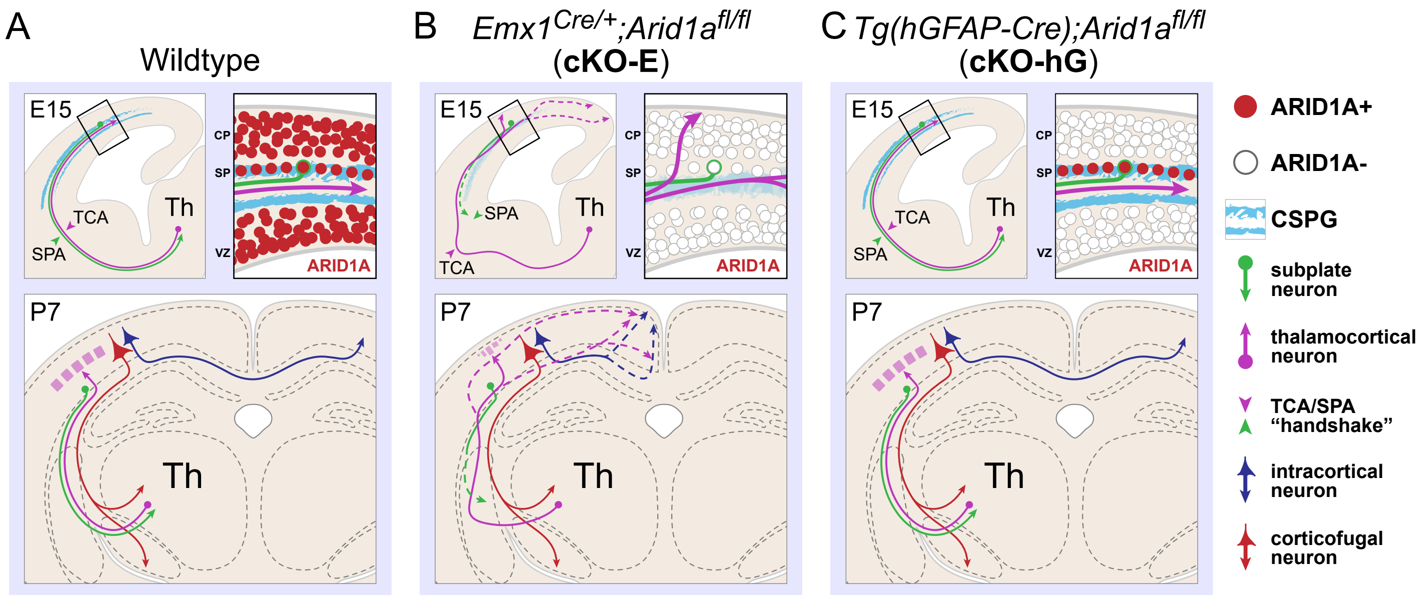


**Fig. S8.** Schematic summary of *Arid1a* function in cortical circuit wiring (*A*) During embryonic cortical development, *Arid1a* is ubiquitously expressed. Descending subplate axons (SPA, green) co-fasciculate with ascending thalamocortical axons (TCA, magenta), a "handshake" interaction proposed to be crucial for reciprocal connectivity between cortex and thalamus (Th). Subplate neurons secrete extracellular matrix (ECM) components, including CSPG (cyan), which form a corridor for the pathfinding of efferent and afferent axons. Upon reaching the subplate, TCAs undergo a "waiting period" prior to invading cortical plate. During the first postnatal week, TCAs extend into cortical layer 4 and form whisker barrels in somatosensory cortex. (*B*) Following pan-cortical *Arid1a* deletion, TCAs (magenta) and callosal axons (blue) are misrouted, and whisker barrel formation is disrupted. Descending subplate axons are attenuated and their co-fasciculation with TCAs is absent. Without the "handshake" interaction, TCAs are impaired in their crossing of the pallial-subpallial boundary. In addition, the subplate CSPG corridor collapses and TCAs, upon reaching the cortex, prematurely invade cortical plate without "waiting" in subplate. Thus, disruption of subplate functions is concomitant with widespread axon misrouting in *Arid1a* cKO-E. (*C*) Following subplate-spared cortical plate deletion of *Arid1a* in cKO-hG, subplate expression of ARID1A is sufficient to support subplate axon co-fasciculation with TCAs and subplate CSPG corridor formation. Remarkably, despite loss of ARID1A from cortical plate neurons, the corpus callosum, and thalamocortical axons and whisker barrels developed without defect. Thus, *Arid1a* plays crucial, non-cell autonomous roles in cortical circuit wiring that are centered on the subplate.

Supplementary Table 1. Genotyping Primers

| Gene Target | Primer_1/Primer_2 (5' – 3') |
|---|---|
| *Cre* | TCGATGCAACGAGTGATGAG |
| | TTCGGCTATACGTAACAGGG |
| *Arid1a*$^{fl}$ | TGGGCAGGAAAGAGTAATGG |
| | AACACCACTTTCCCATAGGC |

Supplementary Table 2. Primary and Secondary Antibodies

| Antibody | Company and product number | Dilution |
|---|---|---|
| Rabbit monoclonal anti-ARID1A | Abcam ab182560 | 1:1000 |
| Rabbit polyclonal anti-TBR1 | Abcam ab31940 | 1:250 |
| Rat polyclonal anti-BCL11B | Abcam ab18465 | 1:500 |
| Rabbit polyclonal anti-LHX2 | Sigma-Aldrich ABE1402 | 1:2000 |
| Rat monoclonal anti-L1CAM | Sigma-Aldrich MAB5272 | 1:1000 |
| Chicken polyclonal anti-GFP | Abcam ab13970 | 1:2000 |
| Rabbit polyclonal anti-GFP | Invitrogen A-11122 | 1:1000 |
| Sheep polyclonal anti-GFP | Bio-Rad 4745-1051 | 1:500 |
| Goat polyclonal anti-Netrin-G1a | R&D Systems AF1166 | 1:100 |
| Chicken polyclonal anti-RBFOX3 | Sigma-Aldrich ABN91 | 1:2000 |
| Rabbit polyclonal anti-CPLX3 | Synaptic Systems 122 302 | 1:1000 |
| Mouse monoclonal anti-TUBB3 | Covance MMS-435P | 1:1000 |
| Rabbit polyclonal anti-NR4A2 | Santa Cruz Biotechnology sc-990 | 1:500 |
| Chicken polyclonal anti-MAP2 | Novus Biologicals NB300-213 | 1:2000 |
| Rabbit polyclonal anti-tRFP | Evrogen AB233 | 1:2000 |
| Rat monoclonal anti-HA | Roche 11867423001 | 1:1000 |
| Mouse monoclonal anti-Chondroitin Sulfate | Sigma-Aldrich C8035 | 1:100 |
| Goat polyclonal anti-SOX2 | Santa Cruz Biotechnology sc-17320 | 1:250 |
| Mouse monoclonal anti-TLE4 | Santa Cruz Biotechnology sc-365406 | 1:250 |
| Guinea Pig polyclonal anti-Cre-recombinase | Synaptic Systems 257 004 | 1:500 |
| Donkey anti-Goat IgG (H+L), Alexa Fluor 488 AffiniPure | Jackson ImmunoResearch Labs 805-545-180 | 1:250 |
| Donkey anti-Rabbit IgG (H+L), Alexa Fluor 488 AffiniPure | Jackson ImmunoResearch Labs 711-545-152 | 1:250 |
| Donkey anti-Rat IgG (H+L), Alexa Fluor 488 AffiniPure | Jackson ImmunoResearch Labs 712-545-150 | 1:250 |
| Donkey anti-Chicken IgY (H+L), Alexa Fluor 488 Affinipure | Jackson ImmunoResearch Labs 703-545-155 | 1:250 |
| Donkey anti-Mouse IgG (H+L), Alexa Fluor 488 Affinipure | Jackson ImmunoResearch Labs 715-545-150 | 1:250 |
| Donkey anti-Mouse IgM, µ chain specific, Alexa Fluor 488 Affinipure | Jackson ImmunoResearch Labs 715-545-020 | 1:250 |
| Donkey anti-Goat IgG (H+L), Alexa Fluor Cy3 AffiniPure | Jackson ImmunoResearch Labs 705-165-147 | 1:250 |
| Donkey anti-Rabbit IgG (H+L), Alexa Fluor Cy3 AffiniPure | Jackson ImmunoResearch Labs 711-165-152 | 1:250 |
| Donkey anti-Rat IgG (H+L), Alexa Fluor Cy3 AffiniPure | Jackson ImmunoResearch Labs 712-165-153 | 1:250 |
| Donkey anti-Chicken IgY (H+L), Alexa Fluor Cy3 Affinipure | Jackson ImmunoResearch Labs 703-165-155 | 1:250 |
| Donkey anti-Rabbit IgG (H+L), Alexa Fluor 647 AffiniPure | Jackson ImmunoResearch Labs 711-605-152 | 1:250 |
| Donkey anti-Chicken IgY (H+L), Alexa Fluor 647 AffiniPure | Jackson ImmunoResearch Labs 703-605-155 | 1:250 |
| Donkey anti-Rat IgG (H+L), Cy5 AffiniPure | Jackson ImmunoResearch Labs 712-175-153 | 1:250 |

Supplementary Table 3. Primers and probes for ddPCR

| Gene Target | Primer_1/Primer_2 | Probe (5' – 3') |
|---|---|---|
| *Ablim1* | ACAGACTTCGCTCAGTACAAC | /56-FAM/ACTACCAGA/ZEN/ |
| | CCTCTGTCCATTCTCACTGC | CCCTCCCAGATGGC/3IABkFQ/ |
| *Myo5b* | GTGACTCTTAACAACCTACTCCTG | /56-FAM/ACAGGCATG/ZEN/ |
| | AAGCCACTCTTCCAGTTGAC | CAACTCAGGTACAACA/3IABFQ/ |
| *Lhx2* | GCATCTGACGTCTTGTCCAC | /56-FAM/TTTCCTGCC/ZEN/ |
| | CTCACCAAGAGAGTCCTCCA | GTAAAAGGTTGCGC/3IABkFQ/ |
| *Tbr1* | CCCGTGTAGATCGTGTCATAG | /56-FAM/TTTAGTTGT/ZEN/ |
| | AGACTCAGTTCATCGCTGTC | GTAATATCCGTGTTCTGGTAGGC/3IABkFQ/ |
| *Tle4* | ACTGACGTGAAAGGAGTATGC | /56-FAM/ACATGCGAG/ZEN/ |
| | AGCCTATGGAAGATCACCTGT | TGCCAGCAATACCT/3IABkFQ/ |
| *Zfpm2* | TGGTTTGTCTGAATGGCTGT | /56-FAM/CATCTGATT/ZEN/ |
| | GAAGACGTGGAGTTCTTTTGTAAC | CTGCTGGCTCCTGGAT/3IABkFQ/ |
| *Srp72* | CTCTCCTCATCATAGTCGTCCT | /5HEX/CCAAGCACT/ZEN/ |
| | CTGAAGGAGCTTTATGGACAAGT | CATCGTAGCGTTCCA/3IABkFQ/ |

### *SI* References


1. O. M. Subach, P. J. Cranfill, M. W. Davidson, V. V. Verkhusha, An enhanced monomeric blue fluorescent protein with the high chemical stability of the chromophore. *PLoS One* **6**, e28674 (2011).
2. T. Matsuda, C. L. Cepko, Electroporation and RNA interference in the rodent retina in vivo and in vitro. *Proc Natl Acad Sci U S A* **101**, 16-22 (2004).
3. J. M. Keil *et al.*, Symmetric neural progenitor divisions require chromatin-mediated homologous recombination DNA repair by Ino80. *Nat Commun* **11**, 3839 (2020).
4. A. Dobin *et al.*, STAR: ultrafast universal RNA-seq aligner. *Bioinformatics* **29**, 15-21 (2013).
5. T. Smith, A. Heger, I. Sudbery, UMI-tools: modeling sequencing errors in Unique Molecular Identifiers to improve quantification accuracy. *Genome Res* **27**, 491-499 (2017).
6. S. Anders, P. T. Pyl, W. Huber, HTSeq-a Python framework to work with high-throughput sequencing data. *Bioinformatics* **31**, 166-169 (2015).
7. M. D. Robinson, D. J. McCarthy, G. K. Smyth, edgeR: a Bioconductor package for differential expression analysis of digital gene expression data. *Bioinformatics* **26**, 139-140 (2010).
8. S. A. Yuzwa *et al.*, Developmental Emergence of Adult Neural Stem Cells as Revealed by Single-Cell Transcriptional Profiling. *Cell Rep* **21**, 3970-3986 (2017).
9. L. Loo *et al.*, Single-cell transcriptomic analysis of mouse neocortical development. *Nat Commun* **10**, 134 (2019).
10. F. Pedregosa *et al.*, Scikit-learn: Machine learning in Python. *the Journal of machine Learning research* **12**, 2825-2830 (2011).
11. F. Wagner, Y. Yan, I. Yanai, K-nearest neighbor smoothing for high-throughput single-cell RNA-Seq data. *bioRxiv*, 217737 (2018).
12. C. R. Harris *et al.*, Array programming with NumPy. *Nature* **585**, 357-362 (2020).
13. M. Waskom, seaborn development team (2020) mwaskom/seaborn. (Zenodo).
14. B. J. Molyneaux *et al.*, DeCoN: genome-wide analysis of in vivo transcriptional dynamics during pyramidal neuron fate selection in neocortex. *Neuron* **85**, 275-288 (2015).
